# 5’ HOXD GENES DIFFERENTIALLY REGULATE GENE EXPRESSION OF SYNOVIAL FIBROBLASTS IN HAND JOINTS

**DOI:** 10.1101/2024.12.10.627720

**Authors:** Masoumeh Mirrahimi, Kerstin Klein, Camino Calvo Cebrián, Alexandra Khmelevskaya, Miranda Houtman, Eva Camarillo Retamosa, Mohammd Hossein Saadat, Ege Ezen, Alexander Vogetseder, Esin Rothenfluh, Martin Berli, Thomas Rauer, Sabrina Catanzaro, Wang Jingyi, Johan Andersson, Oliver Distler, Caroline Ospelt

## Abstract

We previously demonstrated that Homeobox (HOX) transcription factors are differentially expressed between joint locations and can accurately assign synovial fibroblasts (SFs) to their correct joint location. We show here that the expression of the 5’HOXD transcription factors HOXD10, HOXD11, and HOXD13 in SFs strikingly overlaps with predilection sites for the development of rheumatoid arthritis (RA). Changes in SFs gene expression after silencing 5’HOXDs aligned with joint-specific differences of RA SFs. In particular, we identify HOXD13 as regulator or primary cilia function in SFs modulating cell cycle, DNA damage and proteasome activity. Accordingly, we show joint specific differences in primary cilia morphology, DNA damage repair and proteasome activity. We thus propose that HOXD13 and primary cilia play a role in shaping joint-specific SFs functions that might underlie the pathognomic pattern of joint involvement in RA.

## INTRODUCTION

Rheumatoid arthritis (RA) affects approximately 1% of the global population (1) and presents as a symmetric, destructive polyarticular arthritis characterized by a specific distribution of joint involvement, predominantly affecting the small joints of the hands and feet. Notably, the metacarpophalangeal (MCP) and proximal interphalangeal (PIP) joints, particularly the 2 ^nd^ and 3^rd^ MCP and PIP joints, are commonly affected, while the distal interphalangeal (DIP) joints are typically spared. Involvement of the 1^st^ carpometacarpal (CMC) joint is less frequent(2, 3). Though less common, RA can also affect larger joints such as the knees and shoulders, while the hip is less susceptible. Uncovering the local molecular mechanisms specific to each joint is vital for understanding the development of these pathognomic patterns.

Inflammation and degradation in the joints have long been linked to the activation of synovial fibroblasts (SFs) (4, 5). We previously showed that homeobox (HOX) transcription factors and non-coding RNA encoded in the HOX loci are differentially expressed in SF from different joint locations (6, 7). During embryonic development, the formation of joint-specific structures involves precise spatial and temporal regulation of these genes (8). In differentiated cells, little is known about the functional consequences of the preserved site-specific expression. Here, we demonstrate that HOXD10, HOXD11, and HOXD13, encoded in the 5′ end of the HOXD cluster are strongly expressed in distal joints, in particular in digits II-IV, but not digit I. In particular, we identified HOXD13 to be involved in primary cilium formation in SF and to regulate cell cycle, DNA damage repair and proteasome activation. These HOXD13 dependent changes in SF gene expression and function corresponded with differences detected between hand and knee RA SF. Therefore our findings might offer insight into why the small distal joints of the hands are particularly susceptible to developing RA.

## RESULTS

### HOXD10, HOXD11 and HOXD13 are site-specifically expressed in SFs

We first validated the elevated expression of *HOXD10*, *HOXD11*, and *HOXD13 (5’HOXDs)* in cultured SFs from hand joints compared to other sites like elbow, shoulder, and knee, at both mRNA and protein level (Fig. 1a, b). We observed the same distal expression pattern in freshly explanted (Fig. 1c) and in FFPE synovial tissues (Fig. 1d, 1e). During embryonic development a higher dosage of 5’ *Hoxd* in digit II - IV compared to digit I (thumb) shapes the characteristic thumb morphology (9).

**Figure 1:**
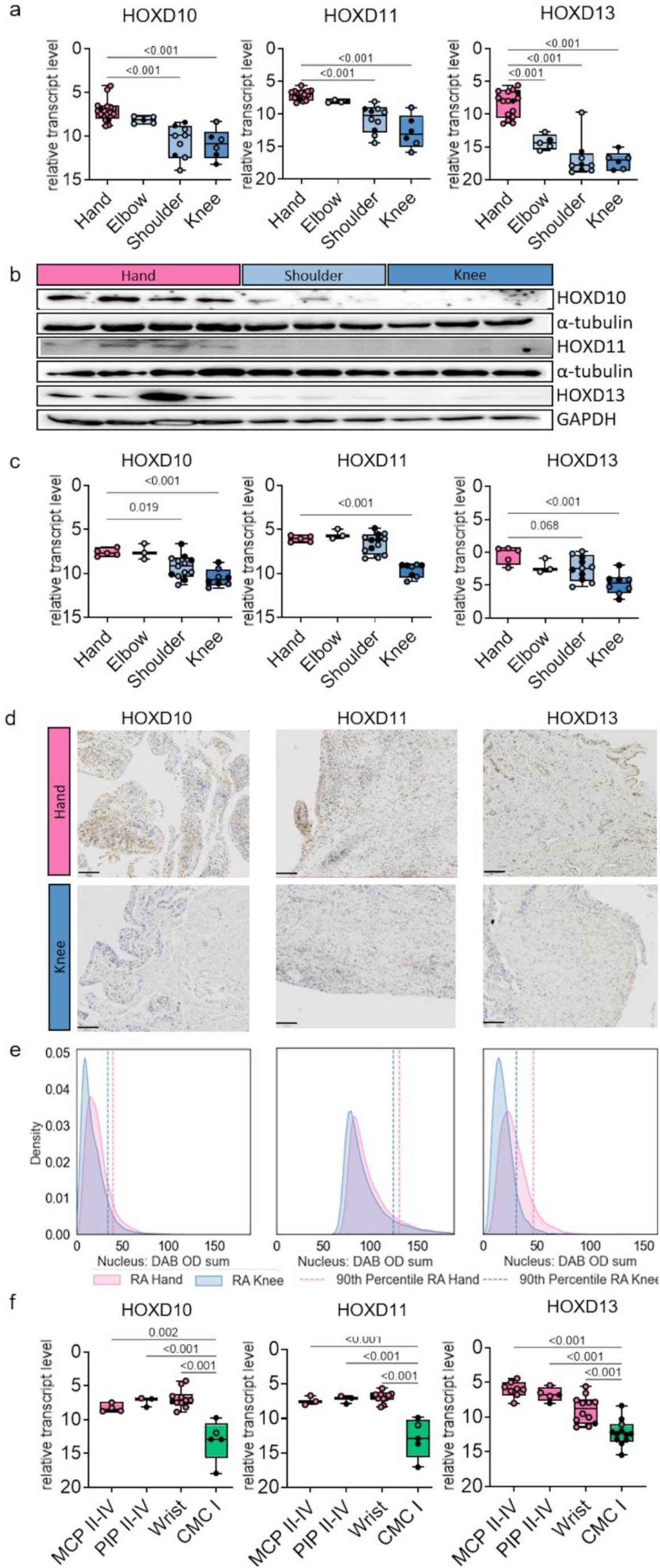
HOXD10, HOXD11 and HOXD13 are site-specifically expressed in SFs. **(a)** Expression levels of *HOXD10*, *HOXD11*, and *HOXD13* measured by qPCR in SFs isolated from hand (n=17), elbow (n=5), shoulder (n=9), and knee (n=6). Ordinary one-way ANOVA with Tukey’s Honestly Significant Difference multiple comparisons test. **(b)** Western blot analysis of HOXD10, HOXD11, and HOXD13 in SFs isolated from hand (n=4), shoulder (n=3), and knee (n=3). **(c)** Expression levels of *HOXD10*, *HOXD11*, and *HOXD13* in synovial tissue isolated from hand (n=5), wrist (n=3), shoulder (n=13), and knee (n=8). Ordinary one-way ANOVA with Tukey’s Honestly Significant Difference multiple comparisons test. **(d)** Representative pictures of synovial tissues from MCP/PIP II-V and knee joints of RA (n=4-6) patients stained for HOXD10, HOXD11 and HOXD13 by immunohistochemistry (IHC). Magnification 200×. **(e)** Quantification of HOXD10, HOXD11, and HOXD13 in RA synovial tissue from MCP/PIP II-V and knee. The figure illustrates the KDE (Kernel Density Estimate) plot for HOXD10, HOXD11 and HOXD13 expression. The x-axis denotes the 3,3’-diaminobenzidine optical density (DAB OD) sum calculated from nucleus of each positive cell, and the y-axis represents the density of samples. The vertical dashed lines indicate the 90th percentile expression levels for each group. Mann-Whitney U test. **(f)** Expression levels of *HOXD10*, *HOXD11*, and *HOXD13* measured by qPCR in SFs isolated from different hand joints: MCP II-V (n=3-8), PIP II-V (n=3-5), wrist (n=10-12), and CMC I (n=5-12). Black filled dots represent OA samples; other dots represent RA samples. Boxplots display the median and range from the 25th to 75th percentile. Whiskers extend from the min to max value. P-values > 0.08 are not reported in the figure.

Accordingly, the expression 5’*HOXDs* was particularly prominent in the MCP and PIP joints of digits II-V and wrists, but not in the CMC joints of digit I (thumb) (Fig. 1f). Similarly, in synovial tissues of the feet, 5’*HOXDs* exhibited lower abundance in the joints of the first digit compared to digits II-V, with a gradually decreasing expression from proximal (metatarsophalangeal, MTP) to distal (DIP) joints (Supplementary Fig. S1a). Consistent with findings in human SFs, 5’Hoxds were lower expressed in DIP joints compared to PIP and MCP joints in C57BL/6 mice (Supplementary Fig. S1b). Taken together, these data show that in SFs the most 5′-located HOX genes, *HOXD10*, *HOXD11* and *HOXD13*, are highest expressed in the more posterior-distal parts of the limbs and that this expression pattern is independent of arthritis and conserved in mice.

### Positional expression of 5’HOXD genes is epigenetically regulated

Analysis of DNA methylation in the 5’ end of HOXD loci revealed higher DNA methylation, particularly in the HOXD10 and HOXD11 regions, in knee-cultured SFs compared to hand-cultured SFs (Fig.2a). Examination of the chromatin landscape in cultured SFs showed that finger SFs exhibited elevated levels of activating H3K27ac marks and lower levels of the repressive H3K27me3 mark compared to shoulder, knee, and thumb SFs. (Fig. 2b,c). Of particular interest were regions, which simultaneously carried both activating (H3K4me3) and low levels of repressing (H3K27me3) marks in finger and thumb SFs, as well as in *HOXD10* and *HOXD11* in shoulder and knee SFs (Fig. 2c and d). Such poised chromatin regions are silent under normal conditions but can be activated by an inflammatory stimulus(10). Indeed, in Assay for Transposase Accessible Chromatin using Sequencing (ATAC-Seq) data, shoulder SFs showed higher accessibility of these genes after TNF stimulation and also knee SFs had increased accessibility of HOXD10 after TNF stimulation (Fig. 2f).

**Figure 2:**
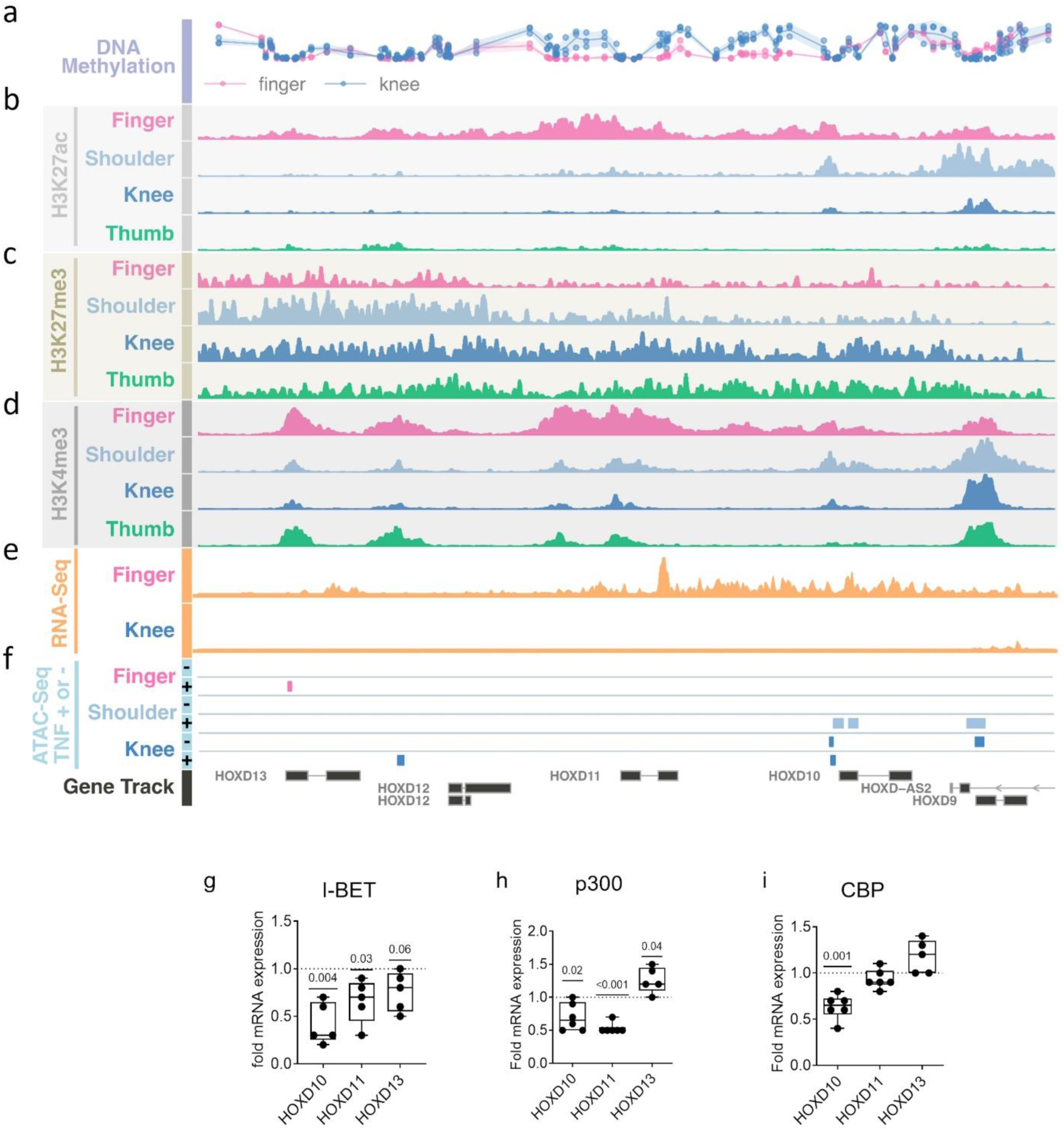
Joint-specific epigenetic regulation of HOXD genes in SFs. **(a)** DNA methylation in RA MCP/PIP II-V SF (n=2) and RA knee SF (n=3) across the 5’ end of *HOXD* loci. Assessment of **(b)** H3K27ac, **(c)** H3K27me3 and **(d)** H3K4me3 marks by ChIP-seq over the promoter/transcription start site region of HOXDs in SFs from RA finger, RA shoulder, RA knee and OA thumb. Data was extracted from GSE163548. **(e)** 5’HOXD genes expression in cultured MCP/PIP II-V (n=3) and knee (n=4) SF from bulk RNA sequencing. **(f)** ATAC-seq data from SF from RA finger (n=2), RA shoulder (n=2) and RA knee (n=2), in unstimulated and TNF-stimulated conditions. Data was extracted from GSE163548. Effects of **(g)** I-BET151, an inhibitor of BET reader proteins, on the expression of *HOXDs* in hand SFs (n=5). Expression of *HOXDs* 48hr after silencing of hand SFs for **(h)** p300 and **(i)** CBP (n=5-6) was measured by qPCR. Boxplots display the median and range from the 25th to 75th percentile. Whiskers extend from the min to max value. One-sample t-test. P-values > 0.08 are not reported in the figure.

We then treated SFs from hand joints with I-BET151, an inhibitor targeting the bromodomain and extra-terminal (BET) protein family members BRD2, BRD3, and BRD4. Proteins within the BET bromodomain family bind to ɛ-N-acetylated lysine of histone 3 (H3) and H4, serving as reader proteins for histone acetylation at actively transcribed sites (11). BET inhibition led to a significant reduction in the expression of *HOXD10* and *HOXD11*, with a weaker effect for *HOXD13* (Fig. 2g). This is consistent with the gradual decrease of H3K27ac marks along the HOXD cluster.

Furthermore, upon silencing of the histone acetyltransferases p300 and CBP (Supplementary Fig. S2b), respectively, there was a significant decrease in the expression of *HOXD10,* but there was no downregulation of *HOXD13*, neither after p300 nor after CBP silencing. Only p300 silencing but not CBP silencing regulated *HOXD11* expression (Fig. 2h, i).

Combined these findings show that the expression pattern of 5’*HOXDs* is epigenetically imprinted in a joint-specific manner by a specific set of histone modifications and epigenetic writers (CBP, p300) and readers (BET proteins). Notably, *HOXD10* and *HOXD11* expression was more reliant on histone acetylation, whereas *HOXD13* was not influenced by modulating reader/writer proteins of this modification. Additionally, TNF stimulation might alter the dynamics of chromatin accessibility in these genes.

### Functional similarity of HOXD10 and HOXD13 in hand SFs

To elucidate the involvement of joint-specific HOXD transcript expression in site-specific gene expression changes of SFs, we silenced HOXD10, HOXD11, and HOXD13, respectively in cultured RA SFs from hand PIP II-V and wrist joints (Supplementary Fig.S3). Silencing of HOXD10 and HOXD13, respectively, regulated a large proportion of overlapping transcripts (40% and 31% of all HOXD10 and HOXD13-regulated transcripts, respectively) (Fig. 3a). Silencing of HOXD11 only overlapped with 18% of HOXD10 -and 16% of HOXD13-regulated genes. Hierarchical clustering on the top 1,500 upregulated and downregulated genes confirmed that the gene expression profiles of after HOXD10 and HOXD13 silencing were more similar to each other than to that of HOXD11 (Fig. 3b). To further investigate functional similarities, we employed Gene Set Enrichment Analysis (GSEA) to uncover significant enriched pathways across the different HOXD target genes. Among the five gene sets used, KEGG and MSigDB Hallmark showed the highest similarities among HOXD10-and HOXD13-regulated pathways, with Jaccard indices of 0.31 and 0.5, respectively (Fig. 3d) (Supplementary Table S1). These results underscore a significant redundancy between HOXD10 and HOXD13, highlighting their closely related functional roles. Part of this overlap might be explained by the downregulation of *HOXD10* after HOXD13 silencing (Supplementary Fig.S3). But also HOXD10 and HOXD11 regulated each other’s expression.

**Figure 3:**
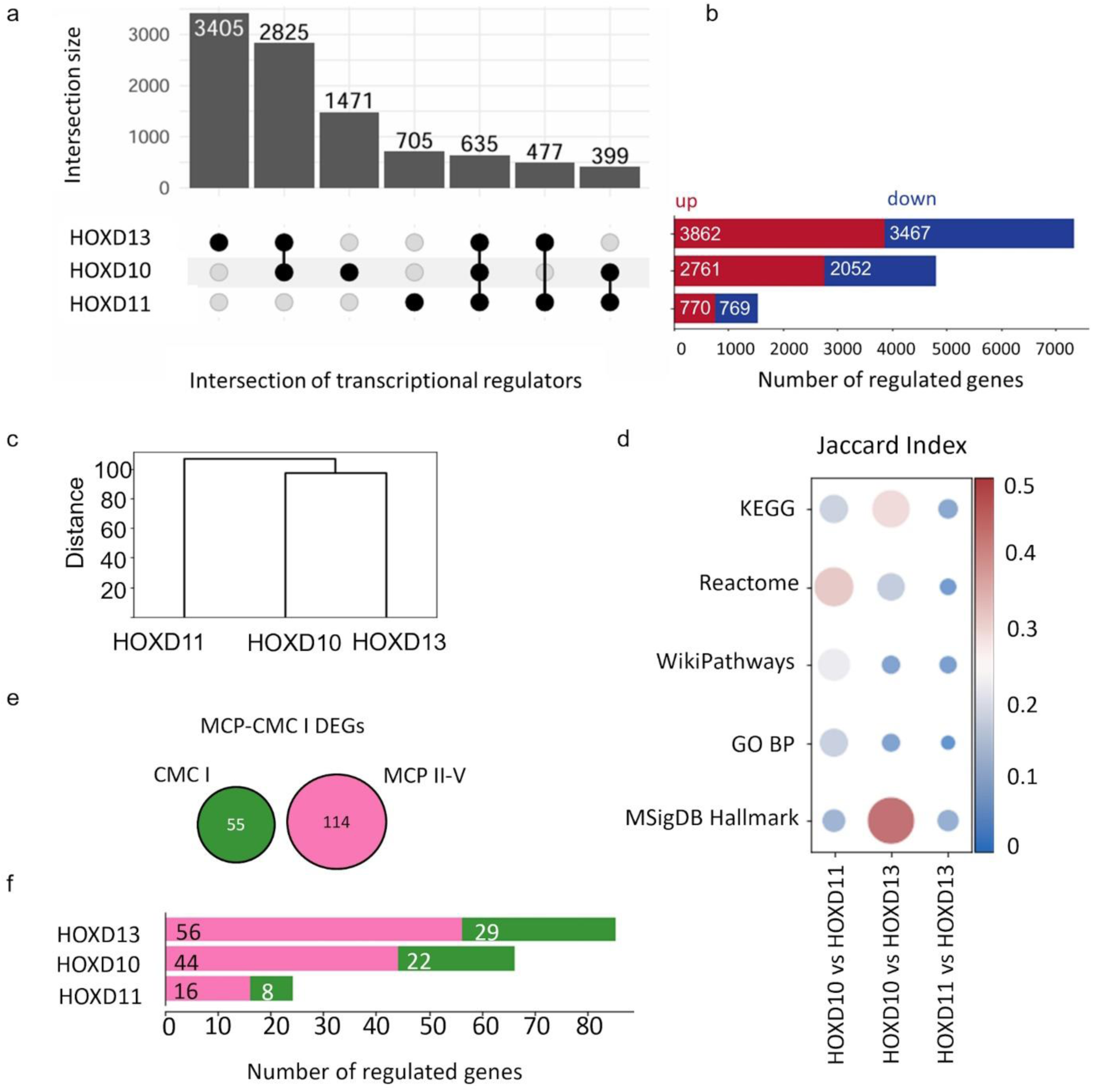
HOXD10 and HOXD13 drive joint-specific gene expression in RA MCP/PIP SFs. **(a)** Overlap between genes altered by HOXDs (10, 11, 13) silencing in SFs transfected with control (GapCTRL) or HOXDs targeting GapmeR after 48hr (Wrist=2, PIP III=1), measured by bulk RNA sequencing. **(b)** Genes up (red) - or downregulated (blue) after HOXDs silencing. |FC | >20% and FDR < 0.05. **(c)** Unsupervised hierarchical clustering of the top DEGs after HOXDs silencing, using 2216 DEGs, which is the size of the smallest cluster (HOXD11). **(d)** Functional similarities were calculated using the Jaccard similarity index on significant pathways (p.adj <0.05) from 5 data bases in GSEA (|FC| >20%, FDR < 0.05). **(e)** Number of genes differentially expressed between MCP II-V (n=3) and CMC I (n=3) SFs in bulk RNA sequencing and **(f)** number of overlaps with genes altered by HOXD10, HOXD11, and HOXD13 silencing. (|FC | >20%, FDR < 0.05).

Additionally, we identified 169 differentially expressed genes (DEGs) between SFs from MCP/PIP II-V and CMC I joints (Fig. 3d). Due to the rarity of RA in thumb joints, we had to use SFs from OA CMC I joints for this comparison. Among MCP-CMC DEGs, 19%, 4%, and 33% were regulated by HOXD10, HOXD11, or HOXD13, respectively, suggesting that HOXD13 in particular plays a role in shaping the hand-specific transcriptome in SFs (Fig. 3e).

### HOXD10 and HOXD13 regulate inflammatory signaling pathways

As suggested by the chromatin data (Fig. 2a-d), the expression of 5’*HOXDs* increased in TNF stimulated SFs and was induced in knee SFs after 48h (Fig. 4a). However, when we overlapped DEGs from bulk RNA-seq analysis of unstimulated and TNF-stimulated cultured MCP II-V versus knee SFs (Fig. 4b), *HOXD10*, *HOXD11*, and *HOXD13* were still the top DEGs after TNF stimulation, highlighting their joint-specific expression under both unstimulated and TNF-stimulated conditions (Fig. 4c). To identify whether 5’HOXDs showed involvement in TNF induced downstream signaling, we compared enriched pathways from HOXDs-silenced DEGs with those from TNF-stimulated MCP DEGs (hand^TNF^) (Supplementary Data 1). Several terms related to TNF, interferon, and the general immune response were negatively enriched in HOXD13 silenced but positively enriched in hand^TNF^ DEGs (Fig. 4d). HOXD10 showed an overlap in TNF signaling and non-inflammatory pathways (Fig. 4d, Supplementary Fig.S3). HOXD11 silencing only overlapped with hand^TNF^ DEGs in pathways unrelated to inflammation (Supplementary Fig.S4). In line with these data, in MCP SFs, a substantial proportion of TNF-regulated genes also were regulated by HOXD13 (Fig. 4e). 47% of downregulated genes after TNF stimulation and 42% of upregulated genes after TNF stimulation were also changed by HOXD13 silencing. Additionally, silencing of HOXD10 overlapped with 47% of downregulated genes and 23% of upregulated genes after TNF stimulation of MCP SFs. The impact of HOXD11 was lower, with only 16% of upregulated genes and less than 1% of downregulated genes being targeted. These findings indicate that HOXD10 and in particular HOXD13, play a crucial role in regulating inflammatory and immune response pathways in SFs from hand joints. Comparison of the expression of 5’HOXDs in synovial tissues from control and RA MCP/PIP II-V indicated higher expression in the context of RA (Fig. 4f).

**Figure 4:**
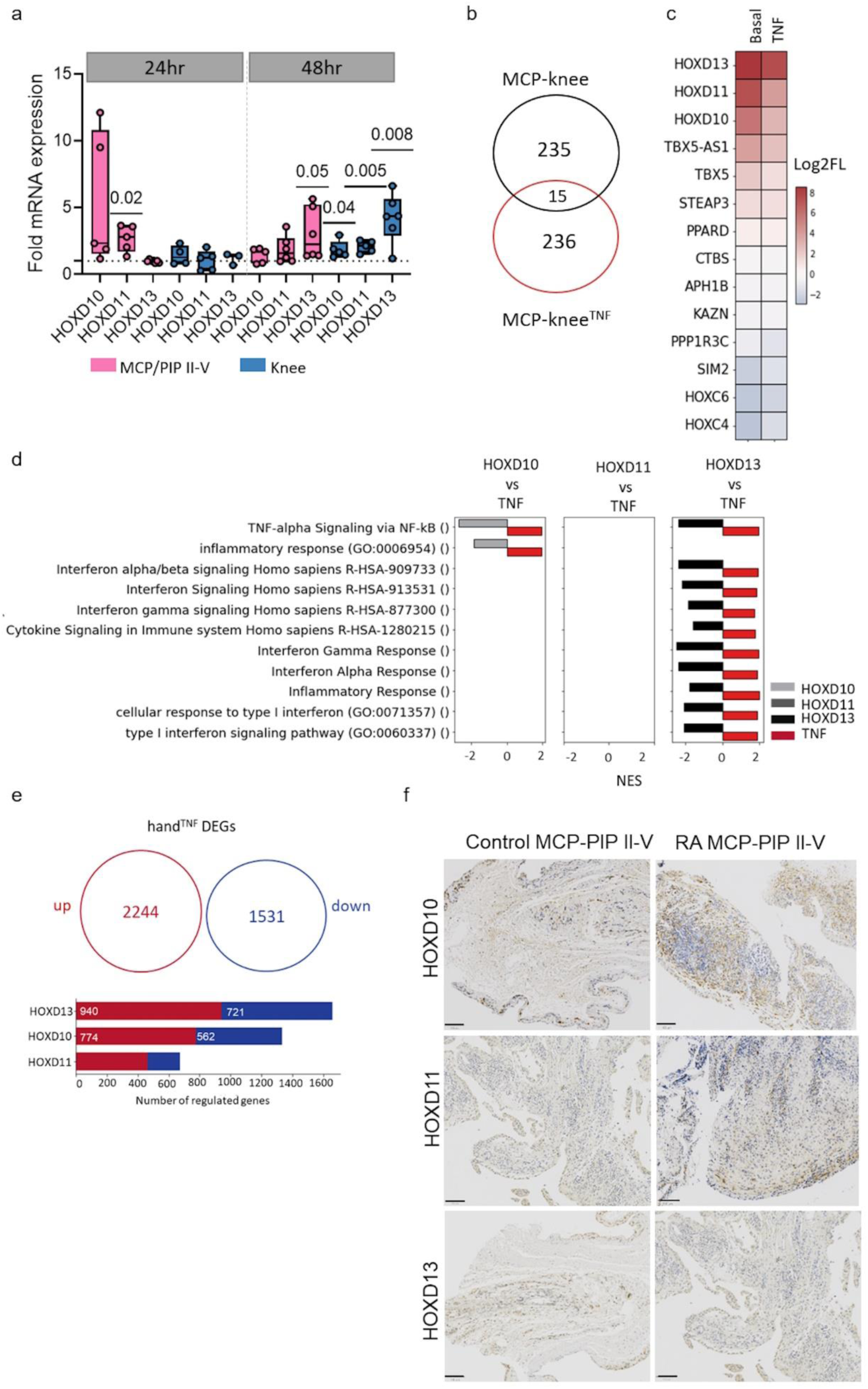
HOXD10 and HOXD13 may regulate inflammatory pathways in MCP/PIP SFs. **(a)** Expression of HOXD10, HOXD11, and HOXD13 were measured by qPCR following TNF stimulation for 24hr and 48hr in SFs form MCP/PIP II-V (n=5-6) and knee (n=5-10). Unstimulated samples were set to 1. Boxplots display the median and range from the 25th to 75th percentile. Whiskers extend from the min to max value. Each dot represents one patient. Statistical analysis was performed using a one-sample t-test. P-values > 0.08 are not reported in the figure. **(b)** Number of all DEGs in hand-knee unstimulated and hand-knee TNF-stimulated (TNF). (| FC | >20%, FDR < 0.05). NES: Normalized enrichment score. **(c)** Heatmap showing the log2 fold change (log2FC) of genes overlapping between hand-knee basal DEGs and hand-knee TNF DEGs. **(d)** Inflammatory-associated terms (p.adj <0.05) commonly enriched in both HOXD^si^ DEGs and hand^TNF^ DEGs. **(e)** Upper panel: number of up-(red) and down regulated (blue) genes after TNF stimulation for 24hr in hand SFs (wrist, n=3; MCP II-V, n=3). Lower panel: number of overlap of these genes with HOXDs (10, 11, 13) DEGs. (|FC | >20%, FDR < 0.05). **(f)** Representative pictures of synovial tissues from RA and control MCP/PIP II-V joints (n=4-6) stained for HOXD10, HOXD11 and HOXD13 by immunohistochemistry (IHC). Magnification 200×.

### HOXD13 regulates cell cycle and DNA damage repair in SFs

We then analyzed the GSEA data comparing 5’HOXD^si^ DEGs with hand^TNF^ DEGs, excluding the inflammation-associated terms. Pathways related to the cell cycle, DNA damage, and apoptosis were the top three pathways commonly enriched (Supplementary Fig. S4, Supplementary Data 1). Further investigation into these pathways showed that most of the terms associated with HOXD10 and HOXD13 silencing were related to the G2/M checkpoint (Fig. 5a) and DNA double-strand break repair (Fig. 6a and Supplementary Data 1). Real-time analysis of SFs revealed no changes in the attachment and spreading, but a significant decrease in the proliferation of HOXD13^si^ cells (Fig. 5b). Similarly, HOXD13 silencing resulted in a reduction in Edu incorporation compared to control transfected SFs (Fig. 5c). No differences in proliferation were seen between MCP/PIP II-V versus knee/shoulder (p=0.45) and MCP/PIP II-V versus CMC I (p=0.88) SFs in basal conditions (Fig. 5d). However, MCP SFs exhibited a significantly higher increase in proliferation compared to knee SFs after stimulation with fibroblast growth factor (FGF) (Fig. 5e). Together these findings suggest that higher expression of HOXD13 in distal joints may contribute to the enhanced proliferative potential of hand SFs.

**Figure 5:**
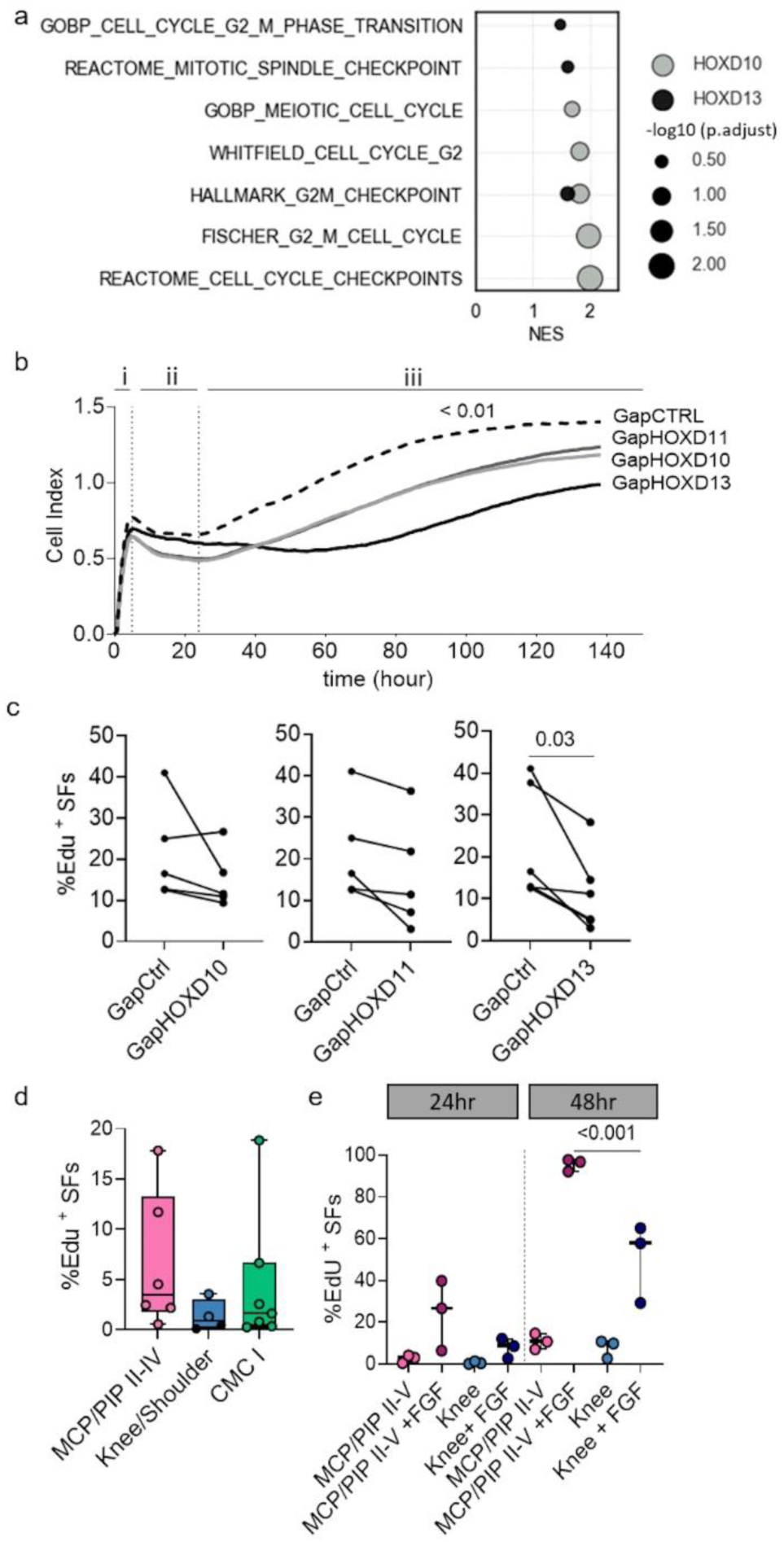
HOXD13 modulates proliferation in MCP/PIP SFs. **(a)** Pathways associated with the cell cycle (p.adj < 0.05) from GSEA on HOXDs^si^ (|FC| > 20%, FDR < 0.05). **(b)** Real-time measurements of i) adhesion (0–5h), ii) spreading (5–24h), and iii) proliferation (24–140h) of RA MCP/PIP II-V SFs transfected with control GapmeR (GapCTRL) or HOXDs GapmeR (n=5). The cell index represents the arbitrary unit of impedance. Data are presented as mean ± standard deviation. Two-way ANOVA. Cell proliferation rate of **(c)** HOXDs^si^ SFs from RA MCP/PIP II-V (n=5) and **(d)** untreated RA MCP/PIP II-V (n=6), RA knee (n=2), RA shoulder (filled black dot, n=2), and OA CMC I (n=7). **(e)** SFs stimulated with bFGF for 24 hours from RA MCP/PIP II-V (n=3) and RA knee (n=3), determined by EdU incorporation after 12 hours of EdU treatment. Boxplots display the median and range from the 25th to 75th percentile. Whiskers extend from the minimum to maximum value. Paired t-test. P-values > 0.08 were not reported in the figure.

**Figure 6:**
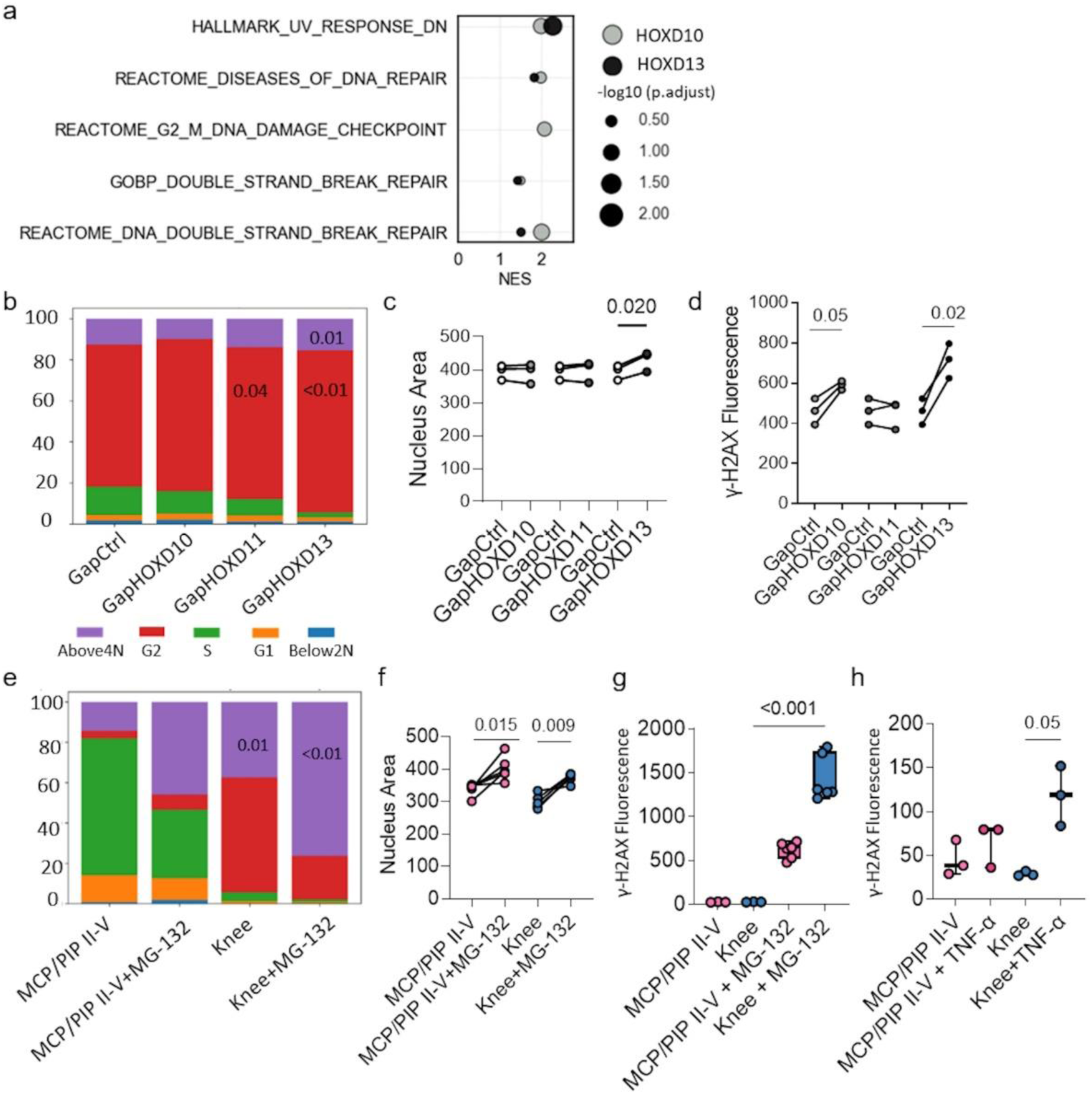
DNA damage response in MCP/PIP SFs is mediated by HOXD10 and HOXD13. **(a)** Pathways associated with DNA damage (p.adj <0.05) from GSEA on HOXDs^si^ DEGs (10, 11, 13). (|FC| >20%, FDR < 0.05). NES: Normalized enrichment score. **(b)** Following silencing of HOXDs, the proportion of RA MCP/PIP II-V SFs (n=3) in different cell cycle phases was assessed. All samples were analyzed in triplicates. Tukey HSD. **(c)** Nucleolus area was measured for each individual cell, and the median for all cells in each patient is represented by each dot in the figure. Paired t-test. **(d)** Quantification of γH2AX foci per cell after HOXDs silencing; each dot represents the median number of γH2AX foci per cell measured for each patient. Boxplots display the median and range from the 25th to 75th percentile. Whiskers extend from the min to max value. All samples were measured in triplicates. Paired t-test. **(e)** Cell cycle phases were assessed in SFs from MCP/PIP II-V (n=5) and knee (n=5) after treatment with MG-132 for 16hr. **(f)** The nucleolus area was measured for each individual cell. **(g)** Quantification of γH2AX foci immunofluorescence staining in untreated and MG-132 treated SFs from MCP/PIP II-V and knee after 16hr (n=3) and 48hr (n=6). **(h)** Quantification of γH2AX foci per cell after TNF stimulation in MCP/PIP II-V (n=3) and knee (n=3) at 48hr.

The intricate regulation of the cell cycle is fundamental to cell proliferation, ensuring that cells divide and replicate in a controlled manner. A significant increase in the proportion of HOXD11^si^ and HOXD13^si^ cells residing in the G2 phase was observed at 48hr (Fig. 6b). Of interest, also the proportion of cells in the “above4N” phase was significantly higher in HOXD13^si^ cells. 4N DNA content refers to the amount of DNA in a cell in the G2/M phase that has duplicated its DNA but has not yet been divided.

The G2/M checkpoint detects and negatively regulates progression through the G2/M transition in response to DNA damage (12). The increase in the G2 phase and in “above4N” in HOXD13^si^ cells suggested a potential halt in progression to mitosis, which was validated by a significant increase in nucleus area after HOXD13 silencing (Fig. 6c). Halting progression to mitosis suggests cell cycle checkpoint impairments and accumulation of DNA damage. Phosphorylation of the histone H2A subtype, H2AX, at Ser139 occurs in response to double-strand break formation, resulting in the formation of γH2AX (13, 14). Following silencing of HOXD10 or HOXD13, cells exhibited increased phosphorylation of H2AX, indicating heightened DNA damage (Fig. 6d).

To actively induce DNA damage, we treated SFs with MG-132 for 16hr (15, 16). MG-132-treated SFs showed a similar increase in proportion of cells in “above4N” phase (Fig. 6e) and increased nucleus area (Fig. 6f). However, while a big proportion of cells went into apoptosis 24hr after MG-132 treatment, silencing of HOXDs for 24hr did not and for 48hr only marginally increased caspase-3/7 activity in SFs (Supplementary Fig. S5).

The DNA damage induced by MG-132 was stronger in knee SFs compared to MCP/PIP SFs (Fig. 6g), suggesting a site-specific response to DNA damage. Additionally, γH2AX levels increased in knee SFs upon TNF-α stimulation, while SFs from MCP/PIP joints remained unaffected (Fig. 6h). Differential analysis of the pathways in MCP-knee^bulk-RNAseq^ DEG indicated that activation of cell cycle checkpoint pathways was significantly higher in MCP/PIP II-V SFs (p value<0.001) (Supplementary Fig. S6a and Supplementary Data 2). Furthermore, the genes involved in the G2/M checkpoint pathway (REACTOME) were higher expressed in MCP SFs in MCP-knee^sc-RNAseq^ DEGs and lower expressed after HOXD10 and HOXD13 silencing (Supplementary Fig. S6b). HOXD11 silencing did not affect the expression of these genes, as shown in the pathway enrichment analysis (Fig. 5a).

These findings underscore that different joints exhibit diverse levels of DNA damage repair, implying specific cellular responses tailored to each joint site. They highlight the critical role of HOXD10 and HOXD13 genes (10, 11, 13) in safeguarding cells against DNA damage by regulating cell cycle progression and DNA damage responses. High expression of HOXDs in SFs might offer protective advantages against DNA harm, particularly in inflammatory and proliferative conditions.

### HOXD13 controls primary cilia biogenesis in a joint-specific manner

Analysis of 5’HOXDs DEGs showed an enrichment of terms related to the assembly and organization of primary cilia in DEGs after HOXD10 as well as HOXD13 silencing (Fig. 7a, Supplementary Data 2). Primary cilia are non-motile, antenna-like structures that play a key role in cell cycle regulation, and have mechano-, chemo-, and osmosensory functions (17, 18). Beyond their role in the cell cycle, primary cilia are involved in several interconnected mechanisms related to DNA damage (19). They consist of nine pairs of microtubules (axoneme) and are predominantly enclosed by the cell membrane. The base of the cilium is connected to the mother centriole, known as the basal body. Primary cilia undergo a series of stages including assembly, elongation, and resorption, which are regulated by the centrosome (20).

**Figure 7:**
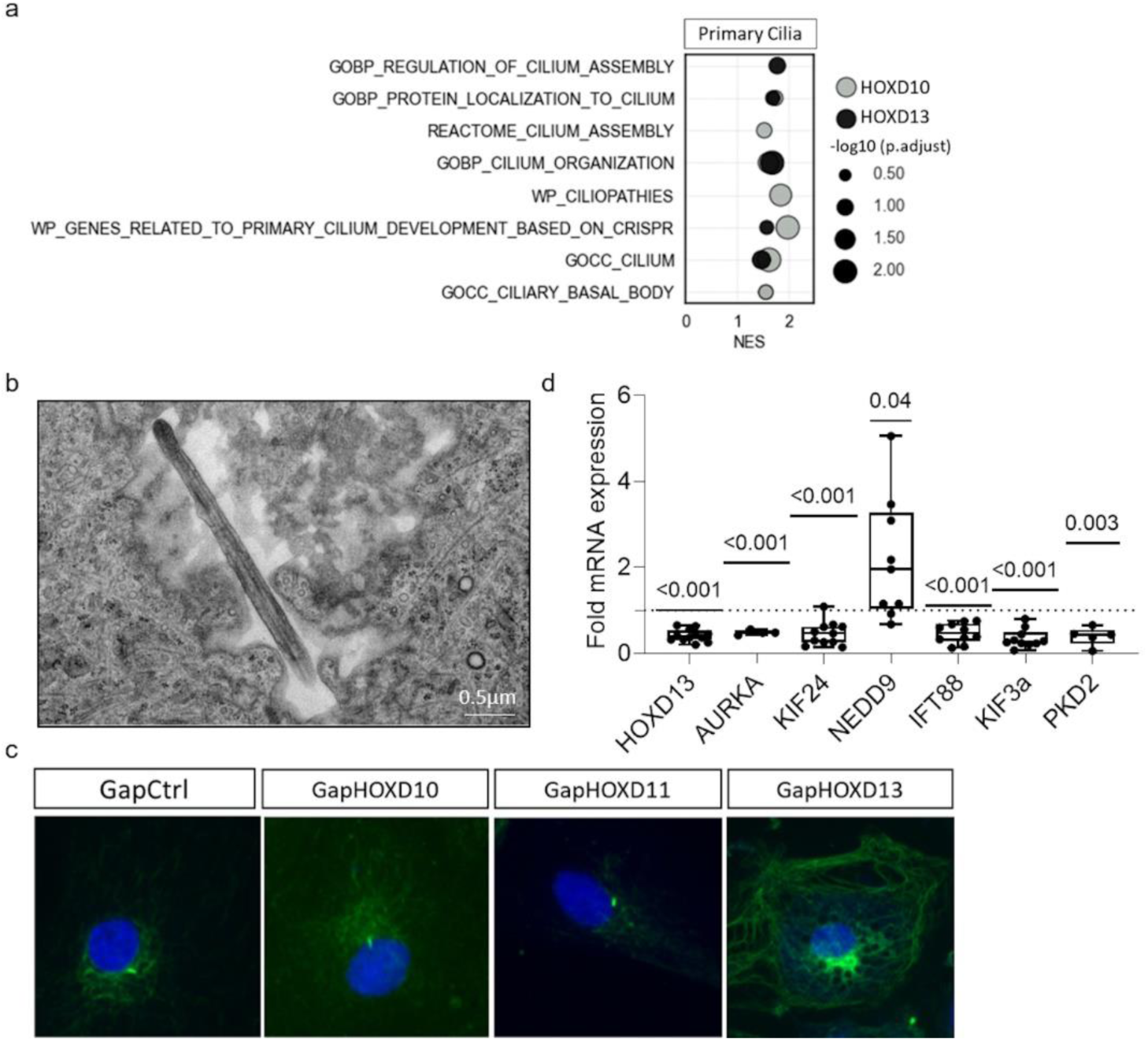
HOXD13 silencing disrupts primary cilia biogenesis in MCP/PIP SFs. **(a)** Primary cilia associated terms (p.adj <0.05) from GSEA on HOXDs^si^ DEGs. (|FC | > 20%, FDR < 0.05). NES: Normalized enrichment score. **(b)** Electron microscopy image showing the primary cilia in SFs. **(c)** Primary cilia were detected using immunostainings of SFs with anti-acetylated alpha tubulin antibodies following silencing of HOXDs. All samples were analyzed in triplicates. **(d)** Expression of genes involved in HDAC6 activity and ciliogenesis after HOXD13 silencing by qPCR (n=6-14). Control transfected cells were set to 1. Boxplots display the median and range from the 25th to 75th percentile. Whiskers extend from the min to max value. Each dot represents one patient. One-sample t test. P-values > 0.08 were not reported in the figure.

Resorption of the primary cilium is necessary to release the centrioles so they can participate in the formation of mitotic spindle poles (21).

We could confirm that SFs harbor primary cilia by electron microscopy (Fig. 7b). Immunostaining with antibodies directed against the cilia marker acetylated α-tubulin clearly visualized the primary cilium in control cells but showed that HOXD13^si^ SFs exhibited hyperacetylation of cytoplasmic microtubules (Fig. 7c). Alpha-tubulin acetylation regulates assembly and disassembly of the axonemal microtubules of primary cilia. Hyperacetylation of cytoplasmic microtubules can indicate impairment in either the de-acetylating enzymes or primary cilia or a combination of both.

The balance of acetylation and deacetylation of microtubules is under the control of ATAT1 and HDAC6 enzymes. The expression of neither of these two genes changed after HOXD13 silencing (Supplementary Fig. S7a), but dysregulation of key factors involved in HDAC6 activity, including *AURKA*, *KIF24,* and *NEDD9*, was observed in RNA sequencing (Supplementary Fig. S7a) (22) and validated by qPCR (Fig. 7d). However, these genes are also involved in cilium disassembly (23, 24).We could confirm previous reports (*25*), that HDAC6 inhibition with tubastatin A (TSA) leads to overexpression of *NEDD9* (26), and downregulation of *AURKA* and *KIF24* (Supplementary Fig. S6b), as seen for HOXD13 silencing (Fig. 7d). However, the two main ciliary genes *IFT88* (27–29), and *KIF3A* (30, 31), were upregulated after treatment with the HDAC6 inhibitor (Supplementary Fig. S7b) and ciliogenesis in SFs was promoted as previously described (22) (Supplementary Fig. S7c). Moreover, after HOXD13 silencing not only regulators of HDAC6 activity, but also several genes with crucial functions in ciliogenesis, including *IFT88* and *KIF3A* were downregulated (Fig. 7d). Therefore, we concluded that the observed hyperacetylation of alpha-tubulin did not result from modulation of HDAC6 activity, but rather from impairments in cilia formation.

To further investigate the involvement of HOXD13 in primary cilia assembly/disassembly we assessed the enrichment of ciliary gene signatures (list of cilia-associated genes obtained from the UniProtKB database, Supplementary Data 2) involved in cilium formation, maintenance, and function (21) in HOXD13^si^ DEGs (Supplementary Fig. S8a). Approximately, 32% of these genes were significantly regulated by HOXD13 (Fig. 8a). These genes could be allocated to various categories based on their roles and positions within the cilia, including ciliary trafficking mediated by the intraflagellar transport (IFT) complex (including the IFT subcomplex, kinesins, dyneins, BBSome), transition zone (TZ), cilium disassembly and ciliogenesis (Fig. 8b). Dynamic changes in the length of primary cilia are under control of the axoneme (20), transition zone (32), IFT complex (33, 34) and BBSome (34–37). The regulation of main genes in each category after HOXD13 silencing pointed towards an overarching role of HOXD13 in maintenance and function of primary cilia.

**Figure 8:**
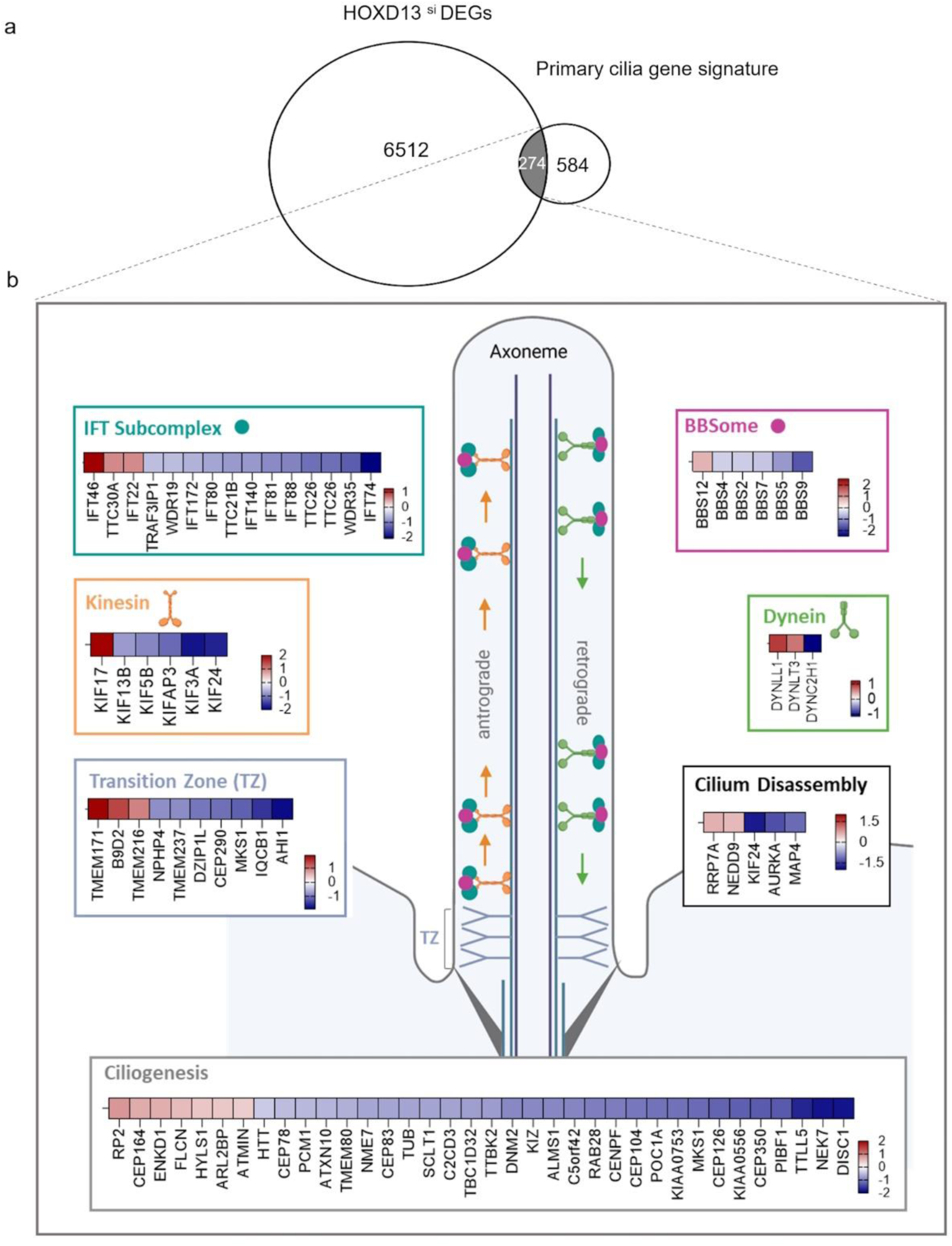
HOXD13 regulates expression of key ciliary genes. **(a)** Overlap of primary cilia gene signature with HOXD13si DEGs (NES=1.3, FDR=0.02). **(b)** Visualization of HOXD13 regulated genes involved into functional categories of primary cilia including intraflagellar transport (IFT) complex (IFT subcomplex, kinesins, dyneins, BBSome), transition zone (TZ), cilium disassembly and ciliogenesis. Red shows up-and blue downregulated genes. Color intensity is proportional to fold change. (|FC| >20%, FDR < 0.05). The image was created using BioRender.

Ciliary associated terms also showed a site-specific enrichment in MCP-knee^bulk-RNAseq^ (Supplementary Fig. S8b and Supplementary Data 2) and MCP-knee^sc-RNAseq^ DEGs (Supplementary Fig. S8c). Serum-starved SFs from different joints exhibited a consistent proportion of ciliated cells, ranging from 75% to 100%, regardless of their anatomical origin (Fig. 9a).

**Figure 9:**
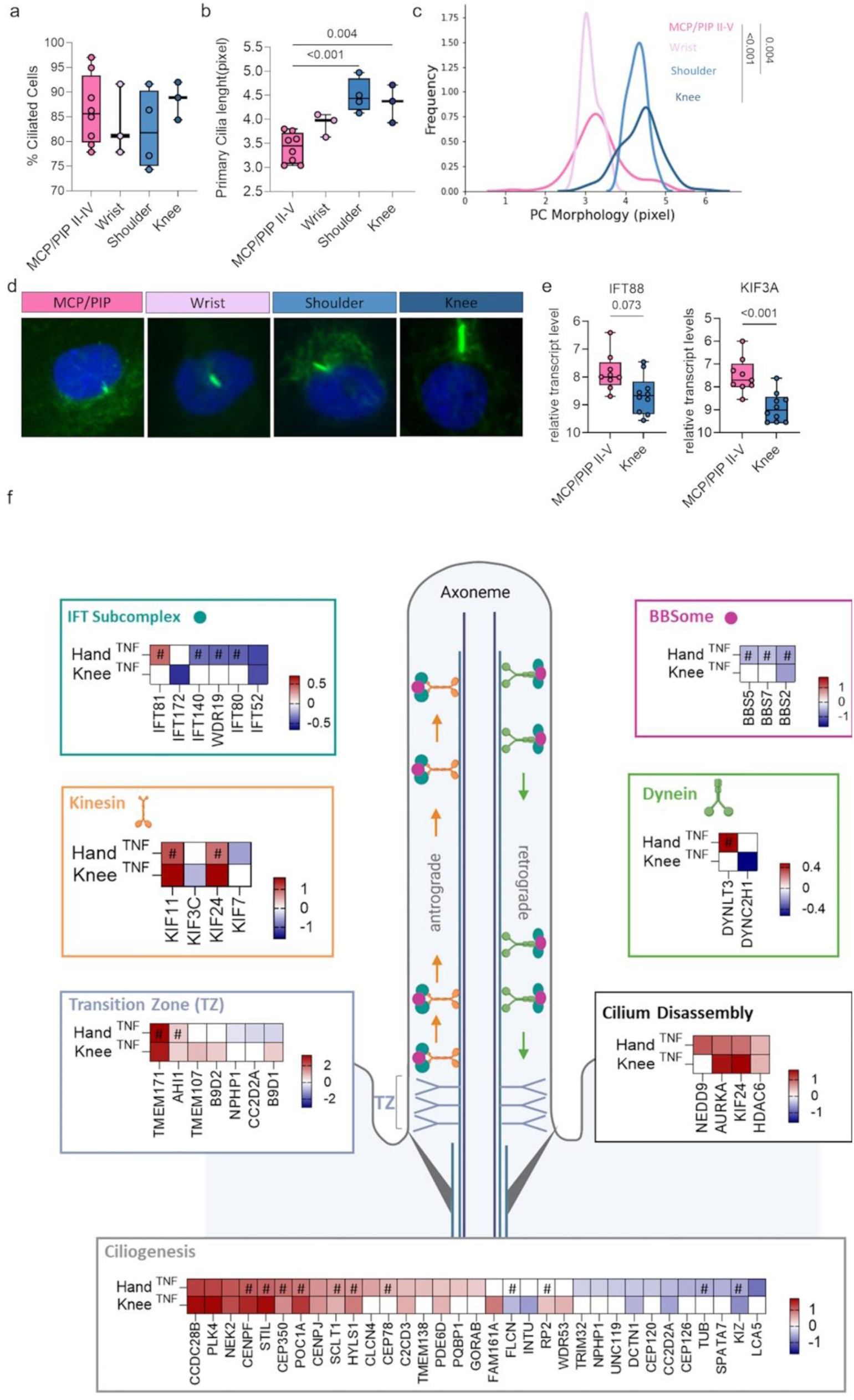
Altered ciliary morphology across different joint locations. **(a)** Fraction of ciliated SFs from MCP/PIP II-V (n=8), wrist (n=3), shoulder (n=4), and knee (n=3). Each dot represents one patient. n=1500-2000 cilia per patient. **(b)** Primary cilia length (two-sided t-test) and **(c)** its distribution in SFs from different joints (two-sided Kolmogorov-Smirnov test). **(d)** Representative immunostaining of primary cilia in SFs using anti-acetylated alpha tubulin antibodies (green). **(e)** Site-specific expression of KIF3A and IFT88 in RA MCP/PIP II-V (n=9) and knee (n=10) SFs, measured by qPCR. One-way ANOVA. **(f)** Visualization of genes involved into functional categories of primary cilia including intraflagellar transport (IFT) complex (IFT subcomplex, kinesins, dyneins, BBSome), transition zone (TZ), cilium disassembly and ciliogenesis that were changed after TNF stimulation of hand (n=6) and knee (n=4) SFs. Red shows up-and blue downregulated genes. Color intensity is proportional to fold change. (|FC| >20%, FDR < 0.05). HOXD13 target genes are indicated by hashtag. The image was created using BioRender. Boxplots display the median and range from the 25th to 75th percentile. Whiskers extend from the min to max value. All samples were in triplicates. P-values > 0.08 were not reported in the figure.

However, primary cilia appeared significantly shorter in serum-starved MCP/PIP II-V and wrist SFs compared to longer ones observed in shoulder and knee SFs (Fig. 9b-d). The site-specific expression of the major genes associated with IFT-and kinesin complexes, *KIF3A* and *IFT88*, was validated by qPCR (Fig. 9e). Upon TNF stimulation, more ciliary genes changed their expression in hand SFs compared to knee SFs, in particular genes involved in the IFT subcomplex and BBSome (Fig. 9f). Of interest, some of these genes were targeted by HOXD13 (Fig. 9f). The genes involved in cilia disassembly were upregulated after TNF stimulation in both locations. This is likely due to the increase in proliferation of SFs after TNF stimulation (38). Given that altered functionality of the IFT machinery and destabilization of the axoneme are hallmarks of primary cilia disassembly, these data suggest that more pronounced primary cilia disassembly in hand compared to knee SFs could be the basis of the shorter primary cilia found in hand SFs.

### Joint-specific modulation of proteasome activity by HOXD13 in SFs

DEGs from HOXD13 silenced SFs showed enrichment for KEGG_UBIQUITIN_MEDIATED_PROTEOLYSIS (NES 1.56, q-value 0.017) (Supplementary Data 2), which is associated with the ubiquitin/proteasome system (UPS). Apart from cell cycle, the formation and resorption of cilia highly depends on UPS proteins, such as E3 ligases and deubiquitinating enzymes. These proteins promote axonemal microtubule assembly and disassembly by regulating the degradation and availability of ciliary regulatory proteins (39, 40). In this context, ubiquitin proteins that localize at the cilium and might regulate cilium formation (41, 42) were among the HOXD13 target genes as well as among the MCP-knee^sc-RNAseq^ DEGs (Fig. 10a). Moreover, a high proportion of proteasome genes (including regulatory 19S subunits and catalytic 20S subunits), known to interact with ciliary proteins (43), were dysregulated by HOXD13 silencing and joint specifically expressed in SFs (Fig. 10a). The relationship between the primary cilium and the UPS system is believed to be reciprocal, as ciliary proteins, in turn, can regulate proteasome activity (44, 45).

**Figure 10:**
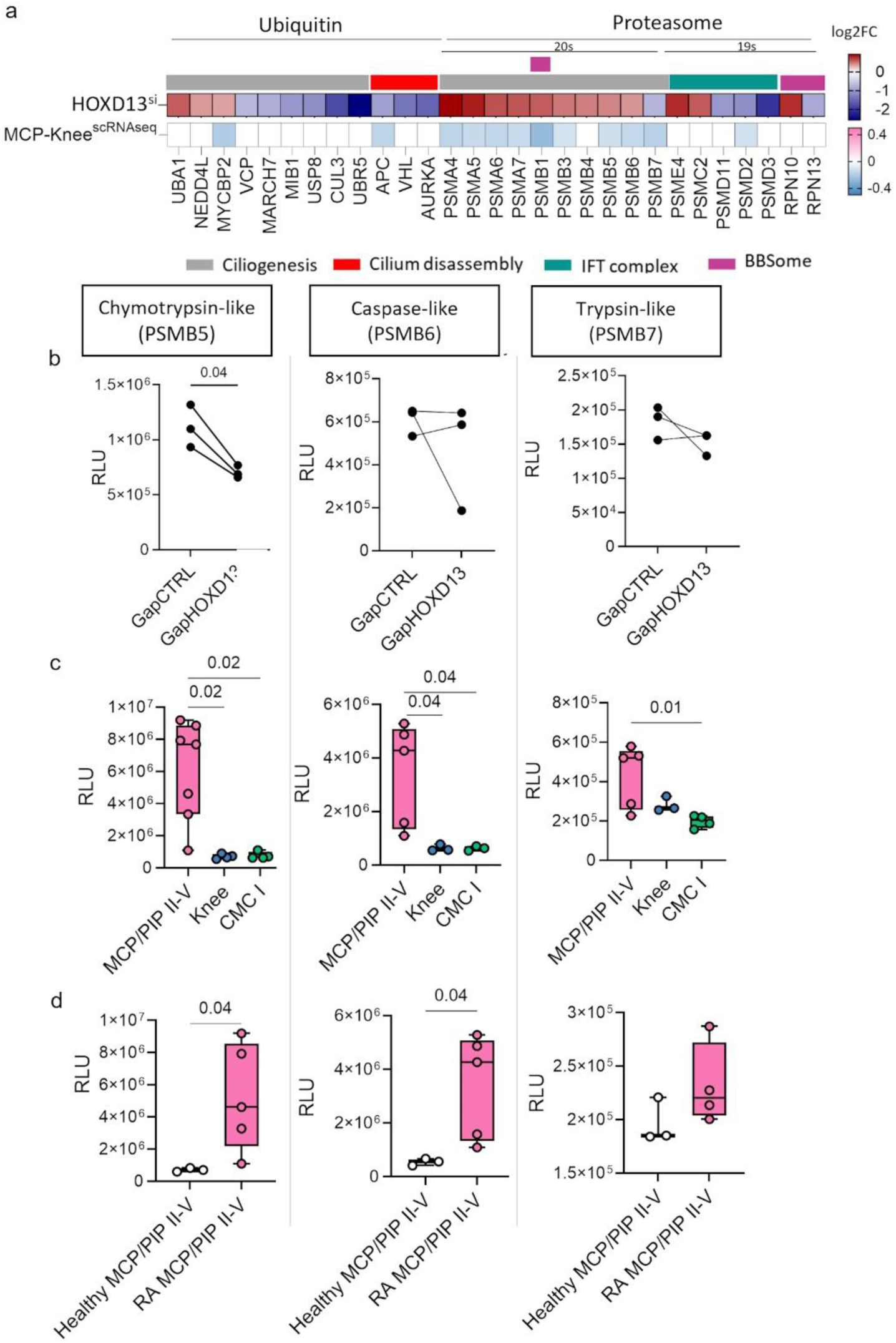
HOXD13 influences joint-specific proteasome activity. **(a)** Heatmap showing log2 transformed fold change (log2FC) for UPS genes associated with primary cilia that were differentially expressed after HOXD13 silencing (n=3, |FC | > 10%, FDR < 0.05) and between MCP (n=3) and knee (n=5) in single-cell RNA sequencing data (|FC | > 20%, FDR < 0.05). Proteasome activity assays performed in **(b)** HOXD13 silenced SFs and **(c)** in RA SFs from MCP/PIP II-V (n=5-7), knee (n=3-4), and CMC I (n=3-4), and **(d)** in SFs from RA (n=5) and control (n=3) MCP/PIP II-V. Data are presented as mean ± standard deviation. Paired t-test (b) and ordinary one-way ANOVA (c, d). P-values > 0.08 were not reported in the figure. RLU = relative luminescence units.

Among the DEGs, there were three subunits of the 20S complex—PSMB5, PSMB6, and PSMB7— which display proteolytic activity equipping the proteasome with chymotrypsin-like, caspase-like, and trypsin-like abilities. We measured these three catalytic activities in HOXD13 silenced MCP/PIP II-V SFs. The chymotrypsin-like activity exhibited a significant decrease following silencing of HOXD13, while there were no significant changes observed in trypsin-like and caspase-like activities (Fig. 10b).

The observed upregulation of 20S subunit genes at the mRNA level might serve as a compensatory mechanism for the reduced enzymatic activities, as previously suggested (46). In SFs isolated from different joints, we measured higher levels of all three activities between cultured MCP/PIP II -V and CMC I SFs and higher activity of chymotrypsin-and caspase-like activity between MCP/PIP II-V and knee SFs (Fig. 10c). Consistent with the negative correlation between transcript expression of proteasome subunits and their protein functions, there was lower expression of 20S subunits in SFs in MCP compared to knee (Fig. 10a), despite higher proteasome activity measured in MCP SFs (Fig. 10c). Chymotrypsin-like and caspase-like proteasome activity was also increased in RA MCP SFs compared to healthy MCP SFs (Fig. 10d). In summary, our data suggests a role for HOXD13 in joint-specific control of the UPS, potentially via stabilization of the primary cilium.

## DISCUSSION

In the present work, we showed that transcripts from the 5′ end of the HOXD cluster (HOXD10, HOXD11, HOXD13) reflect positional identity in SFs. These transcription factors not only distinguish distal (hands, feet) from proximal (shoulder, knee) joints, but also MCP/PIP joints from CMC and DIP joints. This expression pattern strikingly overlaps with joints mostly affected by RA (MCP II-V, PIP II-IV and wrists, thumbs spared), which could indicate a connection between 5’HOXD expression and RA susceptibility.

HOXD13 emerged as a major regulator of site-specific gene expression in hand SF, influencing crucial functions relevant to arthritis. Specifically, we propose that HOXD13 plays a key role in regulating the primary cilium and its associated functions.

In this regard, it is of interest that ciliopathies like the Bardet-Biedl syndrome as well as HOXD13 mutations lead to distal limb malformations such as extra digits (polydactyly), and fusion of digits (syndactyly) (47–49). This overlap in symptoms might reflect a role of HOXD13 in primary cilia structure already during limb formation.

HOXD13 silencing in MCP/PIP II-V SFs led to dysregulation of genes crucial for ciliary trafficking, ciliogenesis, and cell cycle progression. This finding aligned with our cell cycle results, highlighting the role of primary cilia in regulating cell cycle progression and ensuring proper cell division (50–52). HOXD13 silencing impaired cell cycle checkpoints, particularly the G2/M checkpoint, leading to DNA damage accumulation as cells fail to repair damage before progressing through the cycle (53–55). Conversely, it can be assumed that SF with high expression of HOXD13 have more effective DNA damage repair and thus could have a survival advantage, especially under stressful conditions such as chronic inflammation.

In addition to the cell cycle, primary cilia serve as hubs for signal transduction, ensuring that cells respond appropriately to environmental cues (56–58). Ciliary proteins play a crucial role in regulating several cell signaling pathways by local consolidation and by mediating proteasome degradation of signaling mediators and inhibitors (43, 59). Alterations in cilia-regulated proteasome activity can lead to ciliopathies by disrupting key pathways such as Hedgehog (Hh), Wnt, and Notch (59). HOXD13 led to changes in the expression of the ciliary genes BBS2, BBS4, and BBS7, which interact with the proteasome and are involved in Wnt (60) and NF-κB signaling (59). Additionally, we showed that HOXD13 silencing downregulated chymotrypsin-like proteasome activity and measured higher proteasome activity in RA MCP/PIP SFs compared to other joints and compared to healthy samples. Increased proteasome activity has been previously shown in RA SF (61) and might be connected to an improved stress response as well as to the activation of inflammatory signaling pathways via the primary cilium in RA SFs. While numerous studies have demonstrated the role of primary cilia in chondrocytes in OA (62), this study is the first to implicate a role of primary cilia in regulating proteasome and signaling pathways in SFs in RA and to demonstrate a joint-specific architecture of primary cilia. Our finding of shorter primary cilia in MCP/PIP joints points to enhanced signaling through increased local protein concentration (63), as evidenced by higher expression of IFT88 and KIF3A, which are crucial for ciliary trafficking. The length of primary cilia affects the speed of intraflagellar transport (IFT), which is essential for controlling cilia length. This dynamic trafficking is important for the functioning of signaling pathways like Hh, Wnt, and Notch, as it ensures proper localization and movement of signaling molecules within the cilia (64–66). Thus, primary cilia create a microenvironment distinct from the cytosolic compartment, regulating several pathways that were implicated in RA pathogenesis and are considered therapeutic targets (57, 67). Future study might show whether primary cilia may influence RA pathogenesis and could be therapeutically targeted.

The expression of *HOXD10*, *HOXD11*, and *HOXD13* was induced by TNF even in shoulder and knee SFs, despite the presence of repressive H3K27 marks in these genes. This finding aligns with a previous study showing that TNF induces sustained activation of chromatin at regulatory elements associated with genes that evade repression in RA SFs (51). This complex interplay between inflammatory signaling and epigenetic regulation, particularly for genes that are normally expressed in a site-specific manner, suggests that TNF could attenuate these site-specific features at a particular location, potentially disrupting the intrinsic developmental homeostasis of that site. It also provides valuable insights into “joint-specific memory,” a concept describing how certain joints retain unique molecular characteristics that amplify their response to inflammation and disease progression (68).

Our findings confirm and refine our knowledge on differences in SFs function across different joints, but also leave many questions unanswered. The disentangling of differences that occur as a consequence of RA/chronic inflammation and differences that are joint intrinsic would be important for our understanding of joint-specific functions in health and disease. The logistics to sample healthy synovial tissues from various joints are challenging and we could also not obtain RA synovial tissues from joints that are rarely affected by RA such as the CMC I joint. This often left us with comparisons within the disease or across diseases. Furthermore, even though there is a striking overlap of 5’HOXDs expression and RA susceptibility, it will be very challenging to show that the expression of 5’HOXDs indeed promotes the development of RA. The difference between human hand and mouse paw development, morphology and function limit the usage of mouse models to answer this question. Finally and probably most easily addressable is the functional role of the primary cilium in RA. We hope that our results inspire more studies into this intriguing cell organelle and its role in synovial biology, RA and chronic inflammation in general.

## METHODS

### Patients

Synovial tissues were sourced from patients with OA and RA who underwent joint replacement surgery at the Schulthess Clinic Zurich, Switzerland, or from those who underwent ultrasound-guided joint biopsies. RA patients met the 2010 ACR/EULAR (American College of Rheumatology/European League Against Rheumatism) criteria for RA classification, while OA was diagnosed based on chronic pain and evident radiographic signs of OA without underlying inflammatory rheumatic disease. Synovial tissues from feet were sourced from patients who underwent amputation at the Balgrist University Hospital, Switzerland. Healthy knee tissues were obtained using elective arthroscopy after trauma or sports injury at the Clinic for Traumatology, University Hospital Zurich, Switzerland. Healthy MCP and PIP tissues were obtained from patients following hand surgery in which intraarticular procedures were required such as joint fracture reduction, arthrolysis, synovectomy or debridement, at the University Hospital Zurich and the University Hospital Bern, Switzerland. Detailed patient characteristics are provided in (Supplementary Tables S2-S5).

### Culture of SF

#### Human SF

Synovial tissues were digested using Accutase and Liberase at 37°C for 1 hour, and the resulting SFs were cultured in Dulbecco’s Modified Eagle Medium (DMEM; Life Technologies). The culture medium was supplemented with 10% fetal calf serum (FCS), 50 U/ml penicillin/streptomycin, 2 mM L-glutamine, 10 mM HEPES, and 0.2% amphotericin B (all from Life TechnologiesCell cultures were tested and confirmed to be free from mycoplasma contamination using the MycoAlert mycoplasma detection kit (Lonza). SFs from passages 4 to 7 were used for the experiments.

#### Murine tissues

Joints from wild-type C57BL/6 were freed from skin, muscles and ligaments. Whole joints were snap frozen, mechanically crushed and dissociated in 1ml QIAzol using a TissueLyser (Qiagen). For mRNA isolation the miRNeasy kit (Qiagen) with on column DNA digestion was used. The quality and quantity of the RNA was assessed on a NanoDrop machine (ThermoFisher).

### Quantitative real-time polymerase chain reaction (RT-qPCR)

Gene expression was assessed through RT-qPCR. Total RNA extraction was performed using the RNeasy Plus Mini Kit (Qiagen) following the manufacturer’s guidelines, with subsequent cDNA synthesis using the cDNA Synthesis Kit (Thermo Fisher Scientific). FastStart Universal SYBR Green MM (Roche) was utilized for qPCR. Synovial tissue RNA extraction was conducted using the miRNeasy Mini kit (Qiagen) with on-column DNaseI digestion. Control samples, such as no template controls, dissociation curves, and untranscribed RNA samples, were concurrently measured. Data analysis employed the comparative C_T_ method and was represented as 2^−ΔCT^ or 2^−ΔΔCT^. *RPLPO* and *HRPT1* for human SFs and beta2-microglobulin (*B2m*) for mouse tissue were used as internal standards for sample normalization. The list of the primer pairs used in this study is provided for humans in Supplementary Table S6 and for mice in Supplementary Table S7.

### Western blotting

Cells were lysed using Laemmli buffer containing 62.5 mM Tris-HCl, 2% SDS, 10% glycerol, 0.1% bromophenol blue, and 5 mM β-mercaptoethanol. Whole cell lysates were then separated into 10% SDS-polyacrylamide gels and transferred onto nitrocellulose membranes (Whatman) through electroblotting. The membranes were blocked for 1 hour with either 5% non-fat milk or 5% BSA in TBS-T (20 mM Tris base, 137 mM sodium chloride, 0.1% Tween-20, pH 7.6), with BSA used for blocking phosphorylated proteins. Following the blocking step, membranes were incubated with primary antibodies overnight at 4°C. Horseradish peroxidase-conjugated secondary antibodies were then applied. Signal detection was performed using ECL Western blotting detection reagents (GE Healthcare) and visualized with the Alpha Imager Software system (Alpha Innotech). Antibodies and dilutions are listed in Supplementary Table S8.

### DNA Methylation

DNA methylation was performed on SFs from female RA patients (MCP/PIP II-V = 2, knee = 3) using the Illumina Infinium 450k and EPIC BeadChips. The data were then analyzed to obtain the beta values of the CpG sites using the minfi package in R.

### Analysis of ATACseq data sets

ATAC-seq data sets were derived from previously generated data sets (GEO repository accession GSE163548) (69). In brief, cultured SFs form RA finger (n = 2), and RA knee (n = 2) was subjected to ATAC-seq using the standard protocol by Buenrostro et al (70) and sequenced on an Illumina HiSeq 4000. PCR duplicates were identified and removed by Picard Tools prior to peak calling using MACS2 (71).

### Analysis of chromatin-immunoprecipitation sequencing (ChIPseq) data sets

ChIP-seq datasets for H3K4me3, H3K27ac, and H3K27me3 in RA finger, OA CMC I, RA shoulder, and RA knee were previously generated and are accessible at the GEO repository under accession number GSE163548 (6). ChIP data pertaining to the mentioned genomic regions *HOXD10*, *HOXD11*, *HOXD13* were extracted from GSE163548 (GEO repositoryUsing deepTools’ bamCoverage function with Reads Per Genomic Content (RPGC) normalization, the BAM files were converted to BigWig files. These were then visualized, along with DNA methylation, ATAC-seq, and RNA-sequencing tracks, using the Gviz R package.

### Gene silencing

Hand SFs were transfected with 50 nM antisense LNA targeting *HOXD10* (Qiagen, Sequence: 5′-TGT CTG CGC TAG GTG G-3′), *HOXD11* (Qiagen, Sequence: 5′-TGC TAG CGA AGT CAG A-3′), *HOXD13* (Qiagen, Sequence: 5′-CAT CAG GAG ACA GTA T-3′), for 24hr and 48hr for RNA and protein extraction respectively. SF was transfected with 12.5 nM antisense LNA GapmeRs *CBP* (Qiagen, Sequence: 5′-GCG GCG ATC CTT TAG A-3′) or *p300* (Qiagen, Sequence: 5′-TAG TCT GGT CCT TCG T-3′) for 24hr. Transfections were carried out using Lipofectamine 2000 (Invitrogen) in accordance with the manufacturer’s instructions. Antisense LNA GapmeR Negative Control A (Cat. No 300610) served as the transfection control. To analyze differentially expressed genes between hand and knee samples (adjusted p-value < 0.05) regulated by the HOX family, we utilized the R packages UpSetR v14 and ComplexUpset v1.3.3 (72).

### RNA sequencing of control SFs and SFs silenced for HOXD10, HOXD11, HOXD13

Total RNA was isolated from SFs silenced for HOXD10, HOXD11, and HOXD13, as well as control SFs (n = 3 for each), 48hr post-transfection using the miRNeasy Mini kit (Qiagen), which includes on-column DNase I digestion. RNA quantity and quality were assessed using the Qubit RNA BR Assay Kit (Life Technologies) and the Agilent 2100 Bioanalyzer (Agilent Technologies, Inc.). RNA-seq libraries were prepared using the Illumina TruSeq Stranded Total RNA protocol and the TruSeq Stranded Total RNA Sample Preparation Kit. The quality and quantity of the libraries were evaluated with the Agilent 2100 Bioanalyzer using a DNA-specific chip and RT-qPCR with Illumina adapter-specific primers on the Roche LightCycler system (Roche Diagnostics). Diluted indexed long RNA-seq libraries (10 nM) were pooled in equal volumes, used for cluster generation with the TruSeq SR Cluster Kit v3-cBot-HS reagents, and sequenced using the TruSeq SBS Kit v3-HS reagents on the Illumina HiSeq 4000 platform. Genomics was performed at the Functional Genomics Center Zurich (FGCZ) of University of Zurich and ETH Zurich. Sequencing data reads were quality-checked with FastQC, trimmed with Trimmomatic, and aligned to the reference genome and transcriptome (FASTA and GTF files, Ensembl GRCh37) using STAR (73). Gene expression was quantified using the R/Bioconductor package Rsubread version 1.22 (74).

### Analysis of RNA seq data from TNF stimulation SFs

TNF-stimulated SFs RNA sequencing data sets were derived from previously generated data sets available at the GEO repository (accession GSE163548) (69). In brief, cultured RA SFs from hand (wrist, n = 3; MCP/PIP II-V, n = 3) and knee (n = 4) were treated with 10 ng/ml human recombinant TNF for 24 h or were left untreated. Isolated RNA was sequenced using Illumina HiSeq4000/ Illumina Novaseq 6000. All Fastq-files were mapped to hg19, and sequence reads assigned to genomic features using STAR (73) and featureCounts (75). Differential gene expression analysis was performed with DESeq2 R package (version 1.28.1) according to standard protocol (76). Heatmaps were generated using Seaborn library in Python.

### Sc-RNAseq of hand and knee synovial tissues

Single-cell RNA sequencing (scRNA-seq) data were derived from previously generated data sets available in the Array Express repository (accession code E-MTAB-11791). Ultrasound-guided joint biopsies from RA patients, including samples from the wrist (n = 6), MCP/PIP II-V joints (n = 3), and knee joints (n = 6) were processed as previously described (77). Briefly, the tissues were mechanically minced and enzymatically digested with Liberase TL (Roche). For each patient, 10,000 unsorted synovial cells (with viability >80%) were prepared for single-cell analysis using the Chromium Single Cell 3’ GEM, Library, and Gel Bead Kit v3.1, the Chromium Chip G Single Cell Kit, and the Chromium Controller (all from 10× Genomics). The resulting libraries were sequenced on the Illumina NovaSeq6000 platform to a depth of 20,000 to 70,000 reads per cell. Demultiplexing and alignment to the Ensembl reference build GRCh38.p13, as well as collapsing of unique molecular identifiers (UMIs), were performed using CellRanger (v2.0.2) from 10× Genomics. Further analysis followed the standard Seurat protocol (78). Gene expression profiles between hand and knee samples were compared. Detailed patient characteristics are provided in (Supplementary Table S9).

### Differential gene expression and gene set enrichment analysis

Differentially expressed gene analysis was performed in R (version 4.3.1) using DESeq2 package (version 1.42.0) (76) for both bulk-RNA and scRNA sequencing data. Genes exhibiting |FC |>20% and FDR corrected p<0.05 were considered to be differentially expressed. Gene set enrichment analysis (GSEA) and over-representation analysis were performed using clusterProfiler R package (version 4.8.2) (79, 80) using the pathways from the Molecular Signatures Database (MSigDB) (version 7.5.1) (81). Gene sets were considered significantly enriched if the adjusted p-value was below 0.05. A list of genes related to cilia was retrieved from the UniProtKB database (www.uniprot.org) using the keywords "Cilium biogenesis/degradation" (KW-0970) and "Cilium" (KW-0969). These gene lists were then compared with various differentially expressed genes (DEGs) identified through bulk-RNAseq and sc-RNAseq.

### Real-time cell analysis (RTCA)

For real-time cell adhesion and proliferation analysis of SFs, the xCELLigence RTCA DP Instrument (ACEA Biosciences, Inc.) was utilized. Sixteen-well E-plates were prepped by incubating with 100 μl of DMEM, 10% FCS for 30 minutes at room temperature (RT). Prior to cell addition, the impedance of wells with media alone (background impedance – Rb) was measured. SFs were detached using Accutase (Merck), suspended in DMEM, and seeded at a density of 25,000 cells per well. Changes in cell adhesion and spreading, quantified as alterations in impedance, were monitored every 5 minutes for the initial 12 hours (hr) and subsequently every 15 minutes. The Cell Index (CI) at each time point was calculated as (Rn−Rb)/15, where Rn represents the cell-electrode impedance of the well with cells and Rb is the background impedance. Each condition was evaluated in quadruplicate. Impedance changes were recorded every 15 minutes (0– 24hr) and every 30 minutes (24–140hr). Attachment was analyzed during the first 10hr of the experiment, and proliferation was assessed during the exponential phase of the slopes (70-150hr).

### SF stimulation

SFs were stimulated with 10 ng/ml human recombinant TNF (R&D Systems, 210-TA-020), 10 ng/ml human bFGF (Lubio/Peprotech, 100-18B-250UG), 30uM MG-132 (Sigma, M7449-200UL), 10nM Trichostatin A (Selleck, T1952-200UL) in DMEM with 5% FBS.

### EdU proliferation assay

Cells were cultured on 96-well black-walled transparent bottom plate (4 × 10^3 cells per well) in serum-starved medium containing 1% FBS for 24hr to synchronize them in the G0 phase. They were then released into complete medium for 24hr to progress into the G1 or S phase before undergoing HOXD10, HOXD11, and HOXD13 silencing. SFs were treated with 10 µM 5-ethynyl-2’-deoxyuridine (EdU) for 12hr before fixation 36hr post-silencing. To assess the effect of bFGF stimulation on different SFs, DNA synthesis was induced by bFGF (Lubio/Peprotech18B-250UG; 100-10 ng/ml) for 24hr in G1-synchronized cells. EdU incorporation was detected following the Click-iT Plus EdU Alexa Fluor 488 Imaging Kit manual (Thermo Fisher Scientific, C10420). Nuclei were stained with 1 µg/ml Hoechst 33342 (Thermo Fisher, 62249) in PBS for 15 minutes at room temperature, keeping the samples protected from light. Imaging was performed using the Cellinsight CX7 platform (Thermo Fisher Scientific), which is equipped with a 10 × 0.16 NA objective. The analysis was conducted using HCS Studio 2.0 software (Thermo Fisher Scientific). In channel 1, the fluorescence intensity of nuclear DNA stained with Hoechst dye was measured. An image analysis segmentation algorithm was then used to create a nuclear mask to identify viable cells based on size and shape. This mask was subsequently used to measure EdU fluorescence intensity in channel 2 under a fixed exposure time. The proportion of EdU-positive cells was determined and documented.

### DNA damage assay

Cells were seeded onto a 96-well black-walled transparent bottom plate (4 × 10^3 cells per well) and incubated for 24hr. Subsequently, cells were transfected with GapmeR targeting HOXD10, HOXD11 and HOXD13, and GapmeR control for 48hr. After fixation and permeabilization, cells were labeled with a γH2AX antibody (Merk, 05-636; dilution 1:500) to detect phosphorylated histone H2A. Imaging was performed using the Cellinsight CX7 platform with a 10 × 0.16 NA objective and analyzed using HCS Studio 2.0. Nuclear DNA fluorescence (Hoechst dye) in channel 1 was quantified, and a segmentation algorithm created a nuclear mask to identify viable cells based on object area and shape criteria. The nuclear mask was used to measure γH2AX fluorescence intensity of valid objects in channel 2 with a fixed exposure time. At least 1000 valid objects per well were analyzed. An average fluorescence intensity per cell was reported. DNA damage response to TNF-α (R&D Systems,10 ng/ml) stimulation and MG-132 treatment (Sigma, 30 μM) was measured after 24hr and 16hr, respectively.

### Cell cycle analysis

Cells were seeded onto a 96-well black-walled transparent bottom plate (4 × 10^3 cells per well) and cultured in serum-starved medium containing 1% FBS for 24hr to synchronize them in the G0 phase. Subsequently, they were released into complete medium for 24hr to progress into the G1 or S phase before undergoing HOXD10, HOXD11, and HOXD13 silencing. At 24, 48, and 72hr post-silencing, nuclear DNA was stained with Hoechst. For DNA damage induction, SFs were treated with 30 μM MG-132 for 16hr. Images were captured using the Cellinsight CX7 platform with a 10 × 0.16 NA objective and analyzed using the Cell Cycle bioapplication available in HCS Studio 2.0. The primary function of the Cell Cycle bioapplication is to categorize cells into separate phases of the cell cycle (bellow 2N, 2N phase (G1), 2N-4N phase (S), 4N phase (G2), and above 4N) by measurement of cellular DNA content (Hoechst). The percentage of the cells in each phase was reported.

### Caspase3/7 activation measurements

HOXDs-silenced SFs were incubated with CellEvent Caspase-3/7 for 30 minutes at 37°C. The Apoptotic cells with activated caspase-3/7 were measured using the CX7 platform, and the average fluorescence intensity per cell was reported. For a positive control, cells were incubated with MG132 (Sigma; 30 μM) for 24 hours.

### Scanning (SEM) electron microscopy

For SEM, 2-mm-thick slices from one RA synovial tissue were dehydrated using a series of graded acetone baths. The dehydrated samples were then subjected to critical-point drying using a CPD 030 apparatus (Baltec CNC-und Ingenieurtechnik, Berlin, Germany). Finally, the dried tissue slices were coated with gold using a sputter coater (Balzers Instruments, Balzers, Liechtenstein) and analyzed on a transmission electron microscope (H7000, Hitachi High Technologies Corporation).

### Primary cilia staining and imaging

Cells were seeded onto a 96-well black-walled transparent bottom plate (4 × 10^3 cells per well) and cultured in serum-starved medium containing 1% FBS for 24hr to trigger primary cilium assembly. For HOXDs silencing experiments, cells were then released into complete medium for 24hr before undergoing HOXD10, HOXD11, and HOXD13 silencing. Primary cilia were stained with anti-acetylated tubulin antibodies 48hr after transfection. Cells were fixed in 4% paraformaldehyde (Thermo Fisher, 28908) for 20 min at room temperature, washed twice with PBS and permeabilized with PBS containing 0.1% Triton-100 (Sigma-Aldrich, T9284) for 15 min at room temperature. Then cells were blocked with 10% goat serum (Merck/ Sigma, G9023 -5ML) for 1hr at room temperature and incubated with the primary antibody at 4 °C overnight. Cells were washed twice with PBS and incubated with the secondary antibody for 1hr at room temperature. Nuclei were stained using a 0.5 µg/ml Hoechst-PBS solution for 15 min. Subsequently, images and Z-stacks were acquired by the Cellinsight CX7 platform with a 10 × 0.16 NA objective. Z-stacks were collapsed to maximum intensity projections. Antibodies and dilutions are listed in Supplementary Table S10.13

### Primary cilia analysis strategy

Primary cilia quantification was performed using an image segmentation algorithm, known as the watershed algorithm, implemented in Open Source Computur Vision Librarz (OpenCV)(82). It treats the image as a topographic surface, where the intensity of each pixel represents its height. The algorithm identifies the "watershed lines" that separate different objects by simulating the flooding of the surface from the lowest points (intensity minima) until the water from different basins meets. This algorithm is particularly effective for separating overlapping objects in an image, provides precise boundary detection, making it suitable for identifying fine, elongated structures like primary cilia. Major axis lengths are reported as primary cilia length in the figures. The method used here, which provides automatic detection of primary cilia, enables high-throughput screening and large-scale studies (3,000-5,000 cells per condition in our study), facilitating more consistent and reproducible results where relative comparison matters. In our study, given that cell culture conditions were identical across different joint locations and lacked stimuli to alter primary cilia orientation, we infer from these measurements that observed primary cilia morphology is inherently cell-specific.

### Proteasome activity assay

SFs were transfected as described above. Proteasome activities, including chymotrypsin-like, trypsin-like, and caspase-like activities, were measured 48hr post-transfection using the Proteasome-Glo 3-Substrate Cell-Based Assay System (Promega) following the manufacturer’s instructions.

### Histology and immunocytochemistry and imaging

For all analyses, synovial tissue sections at 3 µm thickness were prepared using a HistoCore Autocut microtome (Leica Biosystems, Switzerland) from FFPE blocks and transferred onto Superfrost® Plus Menzel (Thermo Fisher Scientific, CA, USA) glass slides. Histologic tissue staining was performed with hematoxylin and eosin (H&E) for analysis of morphology using standard protocols and reagents.

Immunohistochemistry (IHC) staining for HOXD10, HOXD11, HOXD13 was performed on a BOND-MAX instrument (Leica Biosystems, Switzerland) using the BOND Polymer Refine Detection kit according to manufacturer’s instructions. Briefly, sections were deparaffinized, followed by heat-induced epitope retrieval (HIER) with citrate-based (pH 6, BOND Epitope Retrieval Solution 2) or ethylenediaminetetraacetic acid (ETDA)-based buffer (pH 9, BOND Epitope Retrieval Solution 2). Next, sections were blocked of unspecific antibody binding, followed by incubation with primary and secondary antibodies. 3,3’-Diaminobenzidine (DAB) was used as an antibody detection chromogen and counterstaining was performed with Mayer’s hematoxylin. A list of reagents, antibodies, and instrument settings are listed in Supplementary Table S11. Whole slide scans of histology and IHC stained synovial tissue sections were semi-automatically imaged with an AxioScan.Z1 (Zeiss, Switzerland) microscope scanner at 20x magnification (Plan-Apochromat 20 x /0.8 M27 objective) with a HXP 120 V light source.

### Immunocytochemistry data analysis strategy

The scanned images were analyzed using QuPath version 0.5.0. The threshold for 3,3’-DAB staining was set to 0.09 to minimize false positives and ensure accurate cell detection. Regions of interest (ROIs) were manually defined to include areas of interest while excluding non-tissue regions. A script was created to eliminate detections of cells near boundaries, effectively minimizing both false positives and false negatives. Within each cell detected, the software calculated the sum of DAB staining intensity specifically within the nucleus. Protein expression levels were quantified as the nucleus DAB sum for each cell, reflecting the amount of target protein present. Data from different biological replicates were combined into a single series. The data were then shuffled to ensure randomness, and each dataset was sampled to match the smallest dataset size, standardizing the comparisons across various categories.

### Statistics and reproducibility

All statistical analyses were performed in R (version 4.3.2), Python (versions 3.11 and 3.12), and GraphPad Prism (version 10). Normality of data distribution was assessed using the Shapiro–Wilk test. Two-group comparisons were conducted using two-tailed unpaired or paired t-tests, Mann-Whitney tests, Wilcoxon matched-pairs tests, one-sample t-tests, or one-sample Wilcoxon tests, as appropriate for the data distribution. For multiple group comparisons, Tukey’s Honestly Significant Difference (HSD) test was used following ANOVA to adjust for multiple comparisons. All statistical tests were performed with a two-sided approach, and p-values < 0.05 were considered statistically significant. Each experiment included at least two technical repeats, and all replicates were biological replicates.

### Study approval

The ethical commission of the Kanton Zurich approved the collection and experimental use of human samples (Swiss ethics numbers: 2019-00674, PB-2016-02014, and 2019-00115). Informed consent was obtained from all patients, and all experiments were conducted following institutional guidelines. Surplus mice from the animal facility of the University of Zurich were used for the isolation of mouse joints.

---

**Supplementary Fig. S1:**
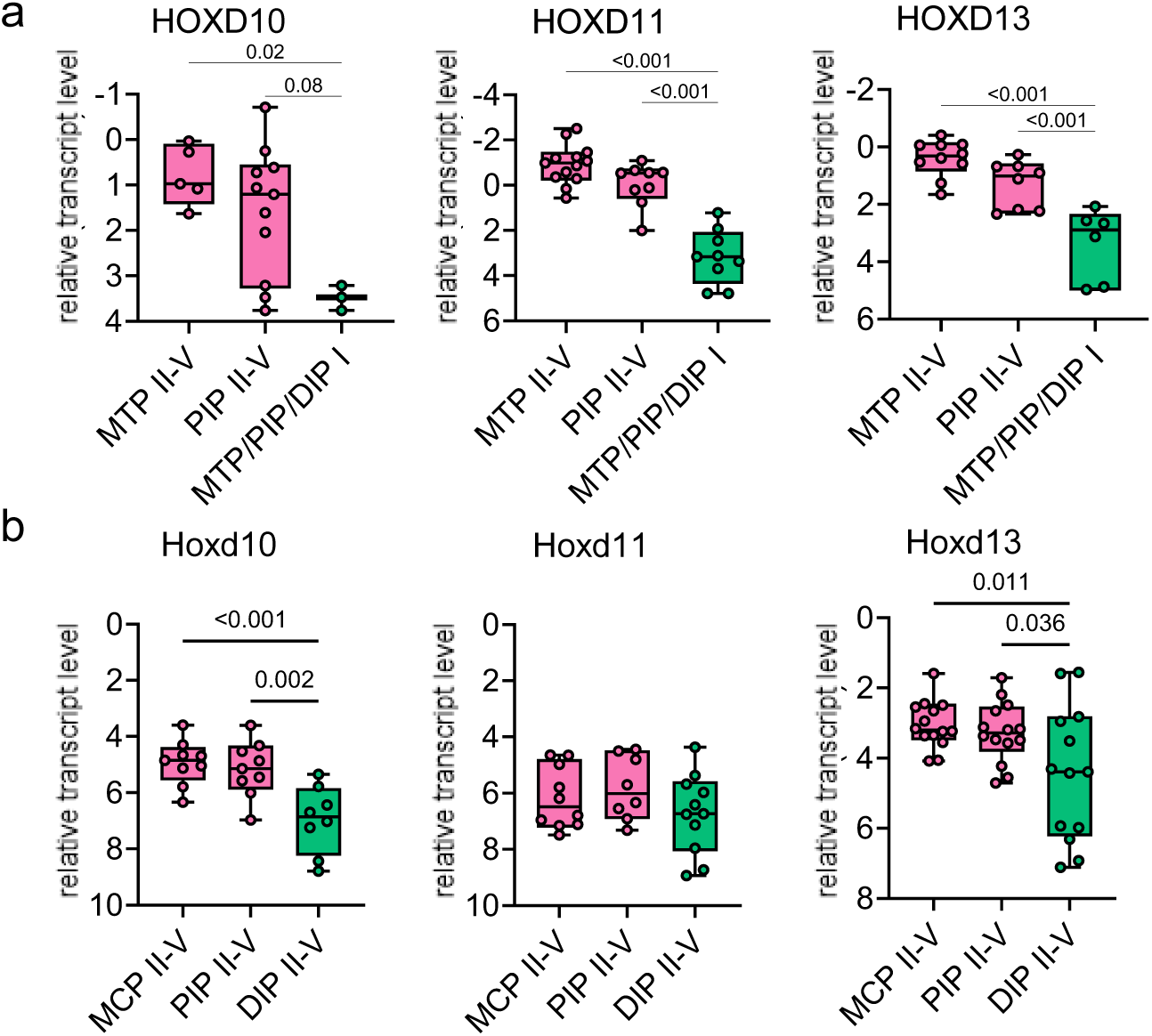
Differential expression of *HOXD10*, *HOXD11* and *HOXD13* in SFs isolated from different joints of human feet and mouse paws. **(a)** HOXDs expression in human feet joints measured by qPCR. MTP II-V (HOXD10 n=5, HOXD11 n=13, HOXD13 n=10), PIP II-V (HOXD10 n=11, HOXD11 n=9, OXD13 n=8), and MTP/PIP/DIP I (HOXD10 n=3, HOXD11 n=9, HOXD13 n=6)). **(b)** HOXDs expression in C57BL/6 mouse paws measured by qPCR. MCP II-V (Hoxd10 8, Hoxd11 n=10, Hoxd13 n=13), PIP II-V (Hoxd10 n=9, Hoxd11 n=8, Hoxd13 n=14), and DIP II-V (Hoxd10 n=8, Hoxd11 n=11, Hoxd13 n=13). Boxplots display the median and range from the 25th to 75th percentile. Whiskers extend from the min to max value. Each dot represents one patient. Ordinary one-way ANOVA. P-values > 0.08 were not reported in the figure

**Supplementary Fig. S2:**
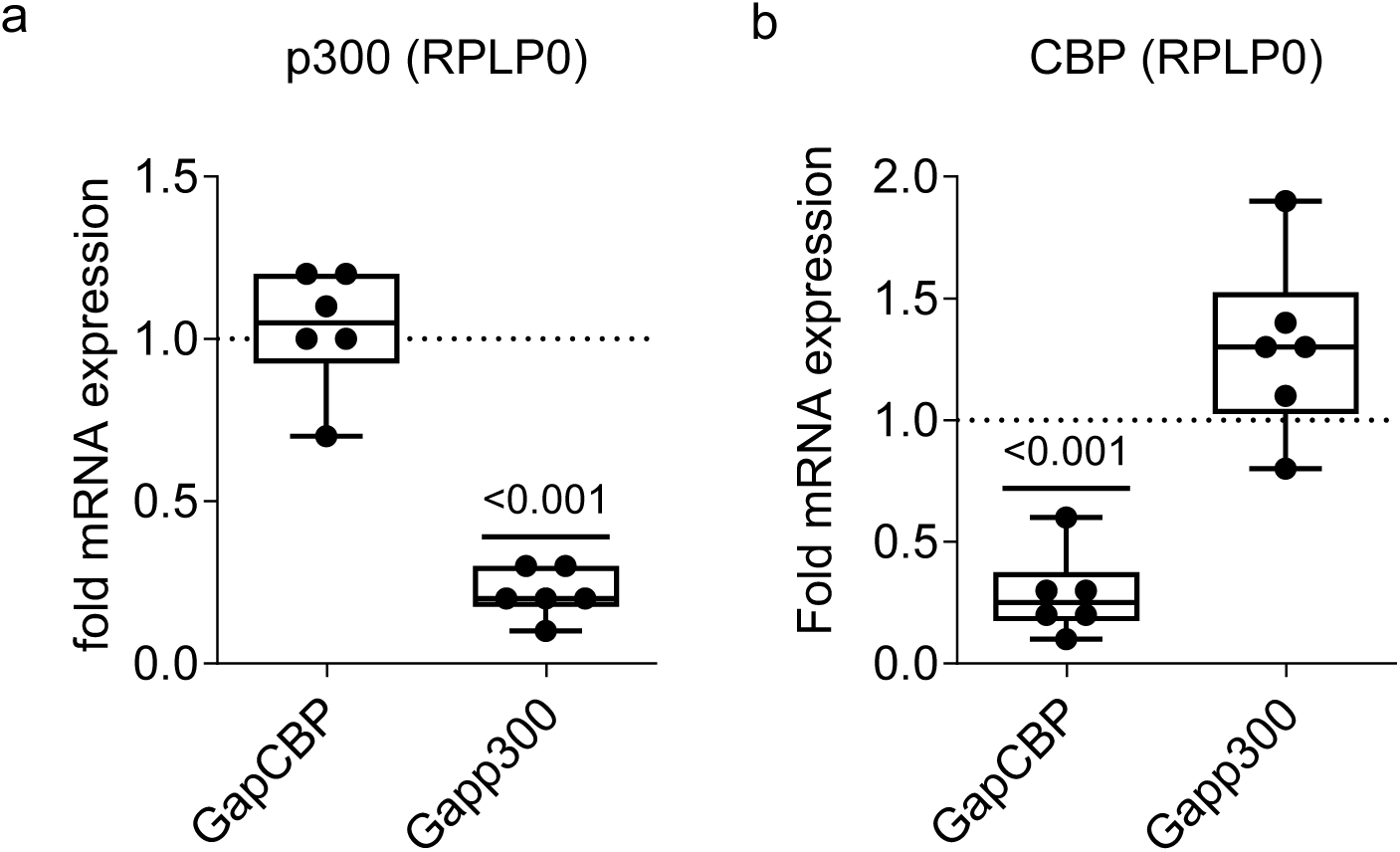
Silencing of CBP and p300 in RA and OA hand SFs. Confirmation of silencing of **(a)** p300 and **(b)** CBP 24hr after GapmeR transfection by qPCR (n=5). Control transfected cells were set to 1. Boxplots display the median and range from the 25th to 75th percentile. Whiskers extend from the min to max value. One sample t test, Wilcoxon signed rank test.

**Supplementary Fig. S3:**
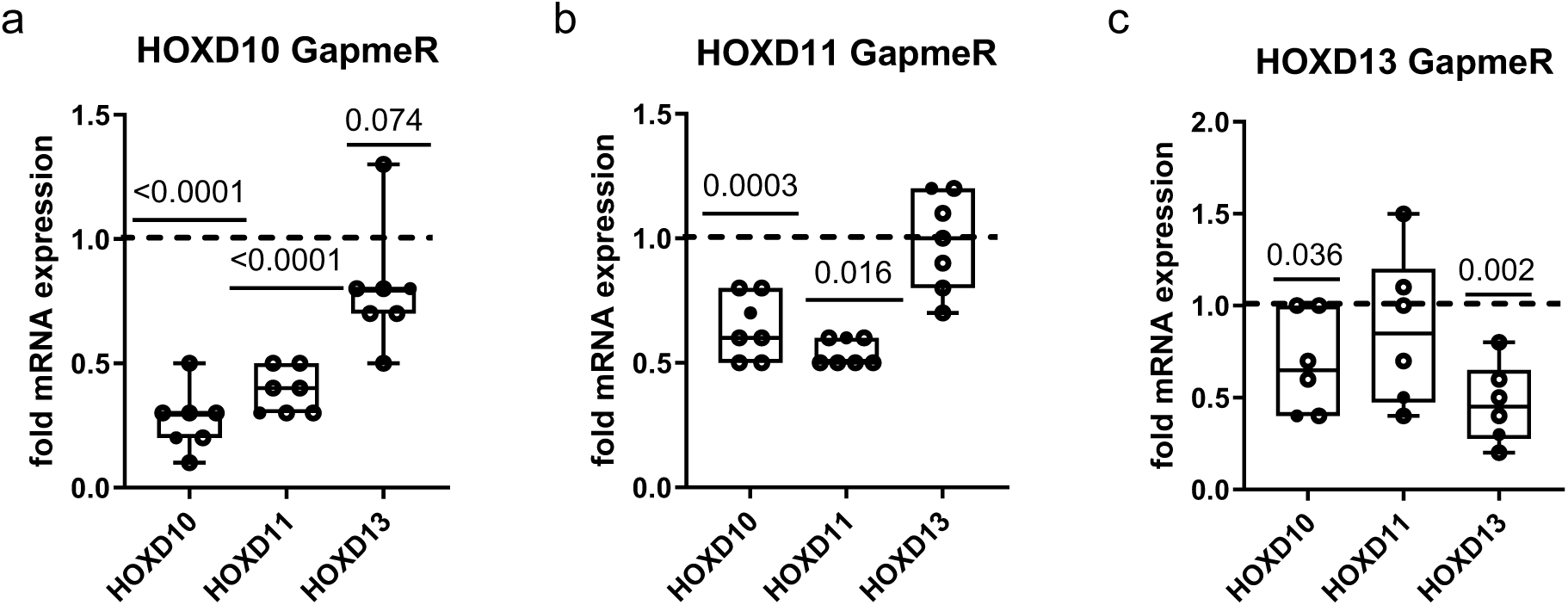
Silencing of HOXD10, HOXD11 and HOXD13 in RA and OA hand SFs. Confirmation of silencing of **(a)** HOXD10, **(b)** HOXD11 and **(c)** HOXD13 24hr after GapmeR transfection by qPCR (n=7). Control transfected cells were set to 1. Boxplots display the median and range from the 25th to 75th percentile. Whiskers extend from the min to max value. All one sample t test, except for HOXD11 in b Wilcoxon signed rank test.

**Supplementary Fig. S4:**
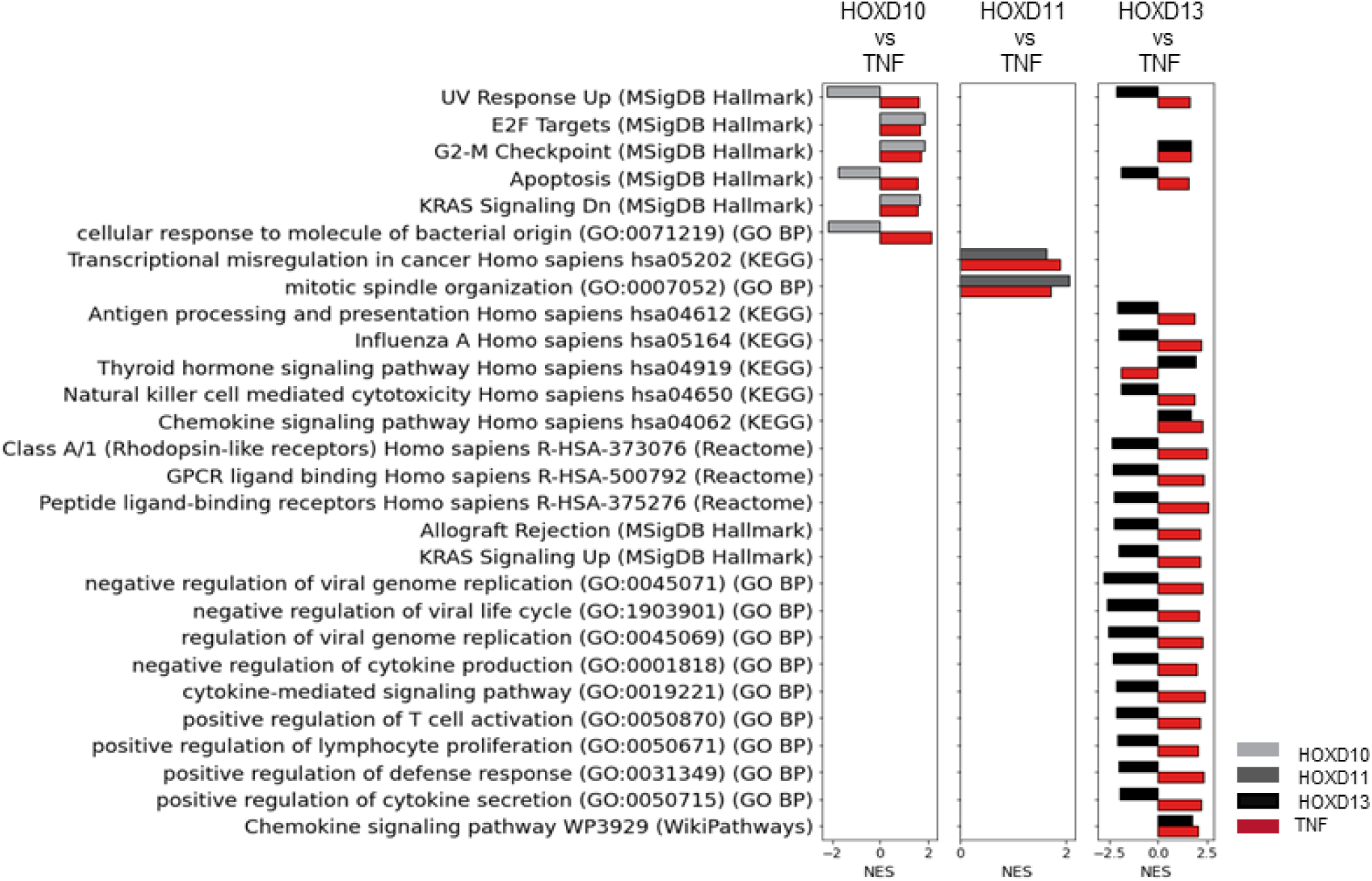
Non-inflammatory terms commonly enriched in both HOXD^si^ DEGs and hand^TNF^ DEGs. Non-inflammatory terms commonly enriched in both HOXD^si^ DEGs and hand^TNF^ DEGs from GSEA across five different gene sets: KEGG, Reactome, Gene Ontology Biological Process, and MSigDB Hallmarks (p.adj < 0.05).

**Supplementary Fig. S5:**
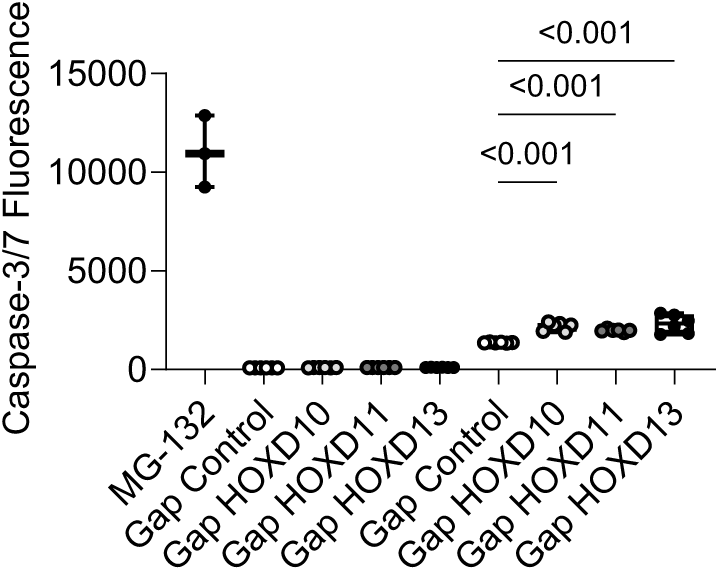
caspase-3/7 activity in SF 24 and 48 hours after HOXDs silencing. Apotosis measurement using caspase-3/7 activity assessment in hand SF (n=6) at 24 and 48 hours post-HOXD silencing, as well as after 24 hours of treatment with MG-132 (n=3). Paired t-test.

**Supplementary Fig. S6:**
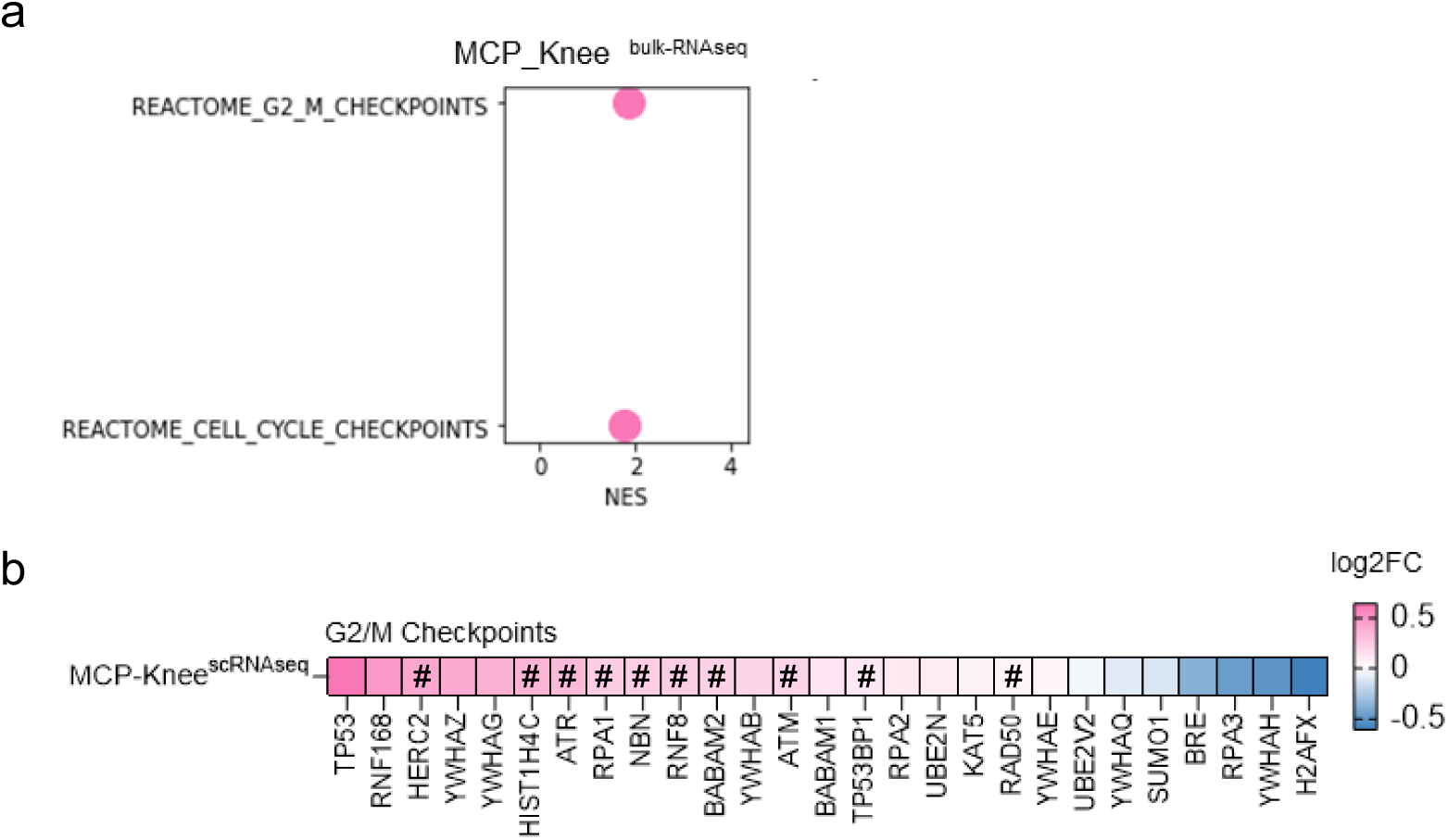
Site-specifc enrichment of cell cycle terms in SFs. **(a)** Enrichment of pathways associated to cell cycle checkpoints in MCP_knee DEGs from bulk RNA sequencing. (p.adj <0.05). **(b)** Heatmap showing the differential expression of genes involved in G2/M checkpoints scRNAseq analysis of MCP and knee SFs. (|FC| >20%, FDR < 0.05). Genes significantly downregulated by HOXD10 and HOXD13 silencing are indicated by a hashtag.

**Supplementary Fig. S7:**
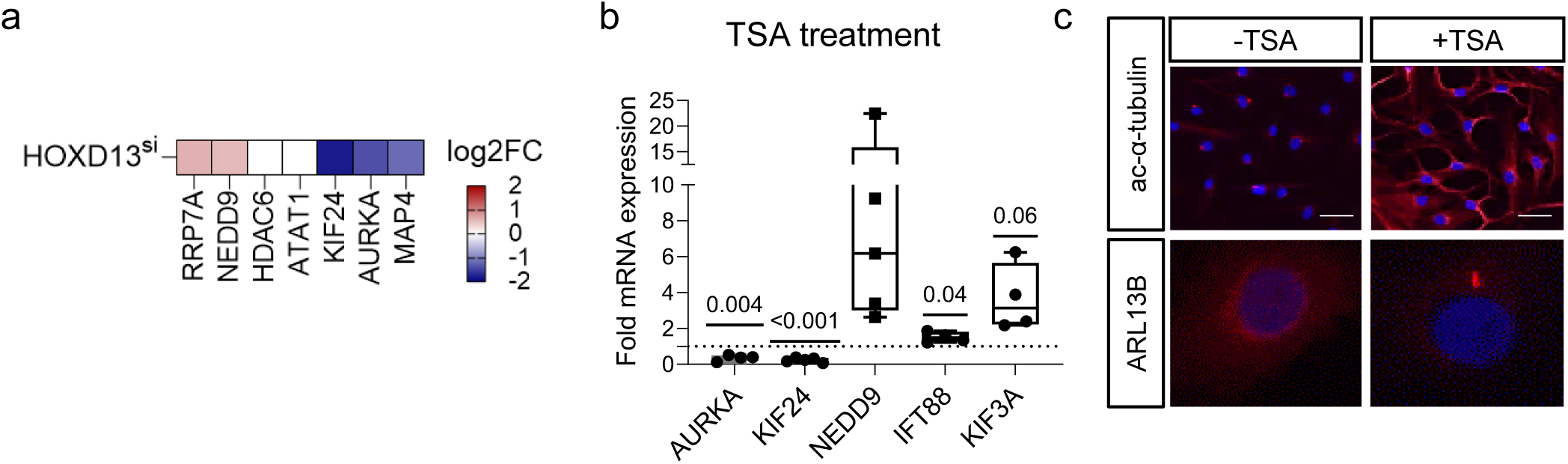
Tubastatin A treatment induced ciliogenesis. **(a)** Heatmap showing the expression of genes involved in HDAC6 activity and cilium disassembly after HOXD13 silencing from RNA sequencing data. **(b)** Expression of key ciliary genes after treatment with the HDAC6 inhibitor tubastatin A (TSA) by qPCR. Boxplots display the median and range from the 25th to 75th percentile. Whiskers extend from the min to max value. Each dot represents one patient. P-values > 0.08 were not reported in the figure. One sample t-test. Control transfected cells were set to 1. **(c)** Representative image of SF treated with TSA. Confirmation of higher acetylation of alpha-tubulin (ac-α-tubulin staining;upper panel) and ciliogenesis (ARL13B staining; lower panel) after treatment. Scale bar= 20um (n=3).

**Supplementary Fig. S8:**
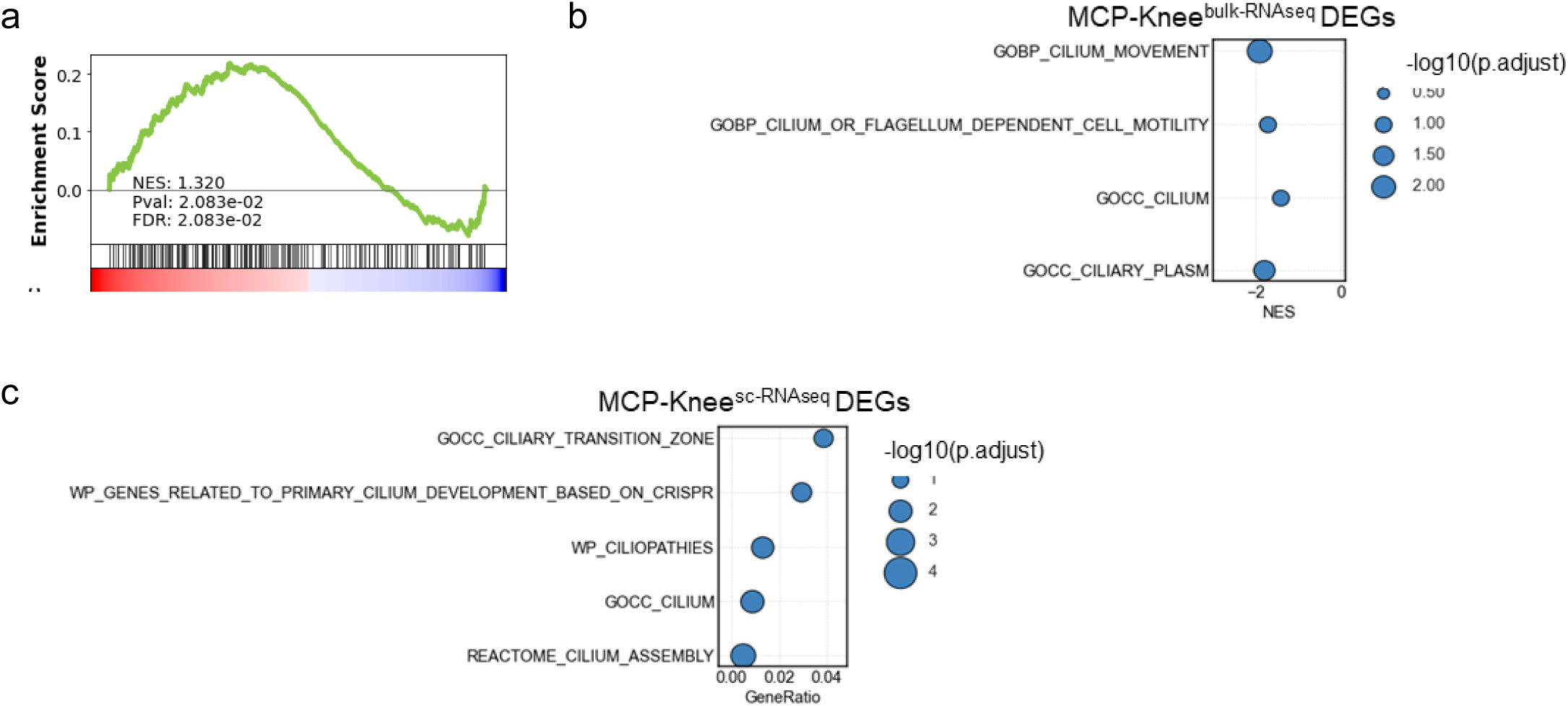
Ciliary genes and pathways exhibit a site-specific pattern in SF. **(a)** Enrichment plot showing significant enrichment of primary cilia gene signature in HOXD13^si^ DEGs. Enriched ciliary associated terms using **(b)** GSEA for MCP-Knee DEGs from bulk RNA sequencing data and **(c)** overrepresentation analysis for MCP-Knee DEGs from single cell RNA sequencing data. (|FC| >20%, FDR < 0.05).

